# Targeted *Tshz3* deletion in corticostriatal circuit components segregates core autistic behaviors

**DOI:** 10.1101/2021.10.15.464549

**Authors:** Xavier Caubit, Paolo Gubellini, Pierre L. Roubertoux, Michèle Carlier, Jordan Molitor, Dorian Chabbert, Mehdi Metwaly, Pascal Salin, Ahmed Fatmi, Yasmine Belaidouni, Lucie Brosse, Lydia Kerkerian-Le Goff, Laurent Fasano

## Abstract

We previously linked *TSHZ3* haploinsufficiency to autism spectrum disorder (ASD) and showed that embryonic or postnatal *Tshz3* deletion in mice results in behavioral traits relevant to the two core domains of ASD, namely social interaction deficits and repetitive behaviors. Here, we provide evidence that cortical projection neurons (CPNs) and striatal cholinergic interneurons (SCINs) are two main and complementary players in the TSHZ3-linked ASD syndrome. We show that in the cerebral cortex, TSHZ3 is expressed in CPNs and in a proportion of GABA interneurons, while not in cholinergic interneurons or glial cells. TSHZ3-expressing cells, which are predominantly SCINs in the striatum, represent a low proportion of neurons in the ascending cholinergic projection system. We then characterized two new conditional knockout (cKO) models generated by crossing *Tshz3*^*flox/flox*^ with *Emx1-Cre* (*Emx1-cKO*) or *Chat-Cre* (*Chat-cKO*) mice to decipher the respective role of CPNs and SCINs. *Emx1-cKO* mice show altered excitatory synaptic transmission onto CPNs and plasticity at corticostriatal synapses, with neither cortical neuron loss nor impaired layer distribution. These animals present social interaction deficits but no repetitive patterns of behavior. *Chat-cKO* mice exhibit no loss of SCINs but changes in the electrophysiological properties of these interneurons, associated with repetitive patterns of behavior without social interaction deficits. Therefore, dysfunction in either CPNs or SCINs segregates with a distinct ASD behavioral trait. These findings provide novel insights onto the implication of the corticostriatal circuitry in ASD by revealing an unexpected neuronal dichotomy in the biological background of the two core behavioral domains of this disorder.

## INTRODUCTION

Autism spectrum disorder (ASD) includes a heterogeneous group of neurodevelopmental pathologies the diagnosis of which is based exclusively on behavioral criteria. The two behavioral domains that are selected by the DSM-5 are: i) deficit in social communication and ii) restrictive, repetitive patterns of behavior, interests, or activities ^1^. These domains also emerge from factor analyses of the 13 available diagnostic instruments in patients ^2^ and in a model that aligns mouse and patient features ^3^. More than 900 genes have been liked to ASD ^4^, among which >100 impact synaptic functions or interact with genes involved in neuronal development ^5^. As a possible neurobiological substrate, clinical and animal studies point to molecular, neurodevelopmental and functional changes of deep-layer cortical projection neurons (CPNs), in particular those of layer 5 (L5) forming the corticostriatal pathway ^6-9^. In this context, we have linked heterozygous *TSHZ3* gene deletion to a syndrome characterized by neurodevelopmental disorders including autistic behavior, cognitive disabilities and language disturbance, with some patients also showing renal tract abnormalities ^10^. *TSHZ3* encodes the highly conserved, zinc-finger homeodomain transcription factor TSHZ3, and has been identified in networks of human neocortical genes highly expressed during late fetal development, which are involved in neurodevelopmental and neuropsychiatric disorders ^9, 10^. It is now ranked as a high-confidence risk gene for ASD (https://gene.sfari.org/database/human-gene/TSHZ3#reports-tab). In human and mouse, high *TSHZ3* gene or protein expression is detectable in the cortex during pre- and postnatal development ^11^. We showed that heterozygous deletion of *Tshz3* (*Tshz3*^*+/lacZ*^) and conditional early postnatal knockout (KO) using the *Camk2a-Cre* promoter (*Camk2a-cKO* mice) lead to ASD-relevant behavioral deficits paralleled by changes in cortical gene expression and corticostriatal synaptic abnormalities ^10, 12^. These data suggest that *Tshz3* plays a crucial role in both pre- and postnatal brain development and functioning, and point to CPNs, and in particular to the corticostriatal pathway, as a main player in the *Tshz3*-linked ASD syndrome. In the mouse striatum, TSHZ3 is not expressed in striatal spiny projection neurons (SSPNs), which represent >90% of striatal neurons, but in a small population of cells that are likely interneurons ^10^. We ^13^ and others ^14, 15^ identified these cells as being mainly striatal cholinergic interneurons (SCINs), whose implication in ASD has been suggested by some studies ^16, 17^. We also showed that the *Camk2a-Cre* transgene is unexpectedly expressed in the SCIN lineage, where it efficiently elicits the deletion of *Tshz3* in *Camk2a-Cre* mice ^13^. Together, these data demonstrate that, within the corticostriatal circuitry, *Tshz3* is deficient in CPNs and in SCINs not only in *Tshz3*^*+/lacZ*^ heterozygous ^10^, but also in *Camk2a-cKO* mice ^12^, which both show the full repertoire of ASD-like behavioral defects. We thus here aimed at investigating the respective contribution of CPNs and SCINs to the pathophysiology of *Tshz3*-linked ASD using targeted conditional deletion of this gene, and provide evidence for the complementary implication of these two neuronal populations in the ASD-related core features.

## RESULTS

### Conditional deletion of *Tshz3* in CPNs

High levels of *Tshz3* gene or TSHZ3 protein expression are detectable in the mouse cortex during pre- and postnatal development ^10, 11^. In the adult cerebral cortex, TSHZ3 is detected in the great majority of CPNs (Caubit et al., 2016). Here, performing immunostaining for beta-galactosidase (ß-Gal) to report the expression of *Tshz3*, we show that *Tshz3* is also expressed in part of cortical GABAergic interneurons, as evidenced using *Tshz3*^*+/lacZ*^*;GAD67-GFP* mice (Fig. S1a). Quantitative analysis indicates that 29.6 ± 1.4% of cortical GAD67-expressing neurons co-express ß-Gal and that these dually stained neurons are rather uniformly distributed among superficial (43.2 ± 2.0%) and deep (56.8 ± 2.0%) cortical layers (n = 12 sections from 2 mice). In contrast, ß-Gal is not detectable in cortical choline acetyltransferase (CHAT) positive neurons (Fig. S1b), Olig2-positive oligodendrocytes (Fig. S1c) and GFAP-positive astrocytes (Fig. S1d, e). To address the role of *Tshz3* in CPNs, *Tshz3*^*flox/flox*^ mice were crossed with *Emx1-Cre* (*empty spiracle homeobox 1*) mice (*Emx1-cKO*). The *Emx1-Cre* mouse expresses the Cre-recombinase in the progenitors of cortical glutamatergic projection neurons (i.e., CPNs) and glial cells from embryonic day 9 (E9), but neither in those of cortical GABAergic neurons, nor of striatal interneurons, including cholinergic ones ^18^. Therefore, in the corticostriatal circuitry of *Emx1-cKO* mice, *Tshz3* function should be specifically lost in CPNs. Compared to control, *Emx1-cKO* mice show a drastic reduction of *Tshz3* mRNA levels and of the density of TSHZ3-positive cells in the cerebral cortex, showing the efficacy of the deletion, while the density of striatal cells expressing TSHZ3 is unchanged (Fig. 1a-c). Despite the loss of *Tshz3* expression in most CPNs, the density of NeuN-positive cells is unchanged (Fig. S2a, b), showing no neuronal loss; in addition, neither the pattern of expression of layer-specific CPN markers, namely CUX1 for L2-4 and BCL11B for L5-6, nor the density of cells expressing these markers is affected (Fig. S2c, d), indicating no major alteration in cortical layering. However, spine density of L5 CPNs from *Thy1-GFP-M;Emx1-cKO* mice is significantly reduced compared to *Thy1-GFP-M* control mice (Fig. 1d, e). By crossing *Emx1-cKO* with *GAD67-GFP* mice, we show that cortical GABAergic neurons still express TSHZ3 (Fig. 1f), confirming the specificity of *Tshz3* deletion in CPNs. To study whether *Tshz3* loss in CPNs could indirectly affect cortical GABAergic interneurons, we compared *GAD67-GFP* control mice (Control-*GAD67-GFP*) to *Emx1-cKO-GAD67-GFP* mutant mice. No significant changes in the number of GABAergic interneurons (Control-*GAD67-GFP*: 140.7 ± 4.9, n = 37 sections from 5 mice; *Emx1-cKO-GAD67-GFP*: 144.6 ± 6.1, n = 41 sections from 7 mice; *P* = 0.624, Student’s *t*-test) and in their distribution are found (Fig. S3a, b). CHAT immunostaining on striatal slices in *Emx1-cKO* mice also shows no significant modification of the density of SCINs (Fig. S3c, d).

**Figure 1.**
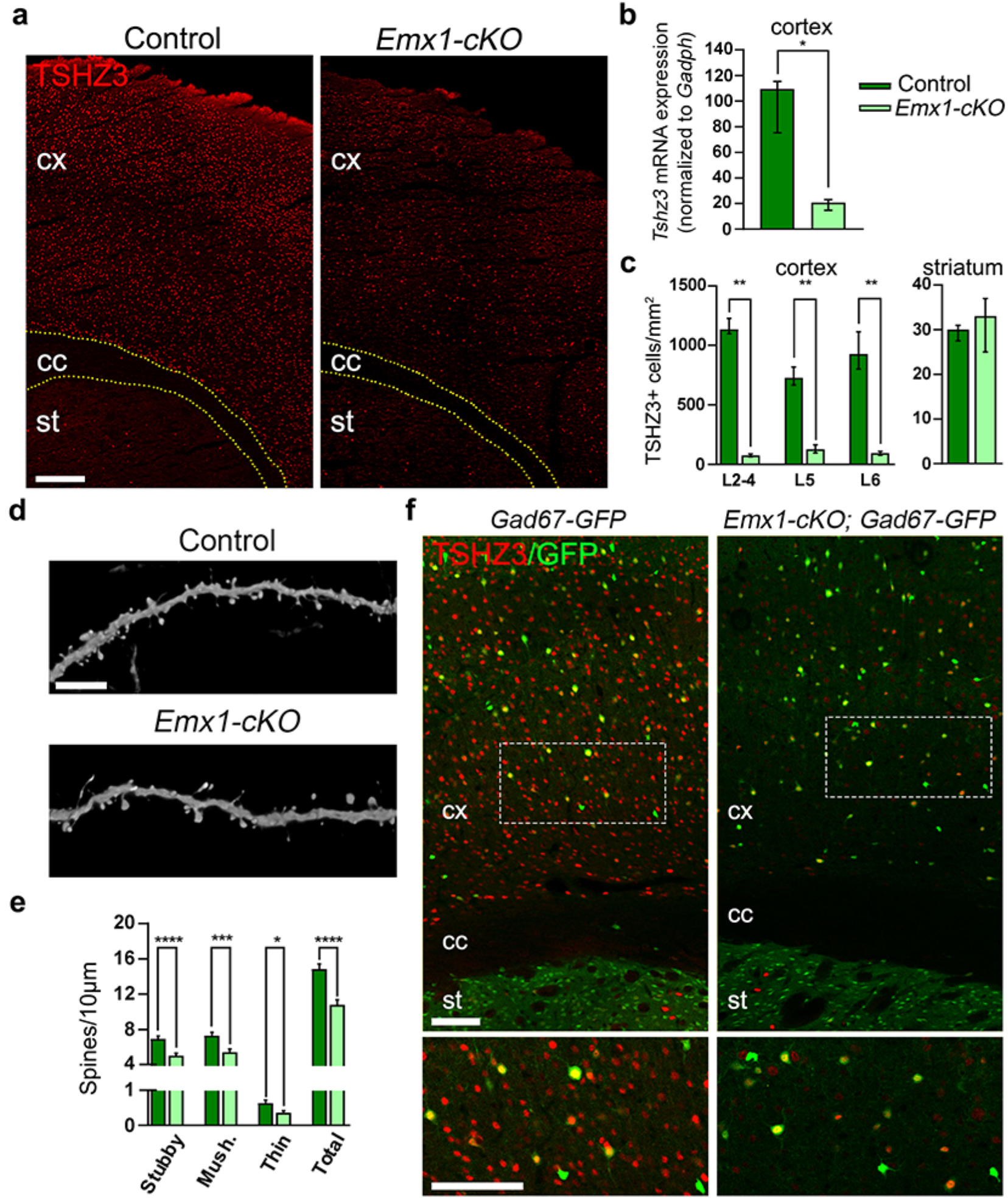
Conditional *Tshz3* deletion in CPNs. **a** Coronal brain sections from control and *Emx1-cKO* mice immunostained for TSHZ3. Scale bar 250 µm. **b** *Tshz3* mRNA relative expression in the cortex of control and *Emx1-cKO* mice measured by RT-qPCR (4 cortices per group; \**P* < 0.05, Mann-Whitney test). **c** TSHZ3-positive cell density in control and *Emx1-cKO* mice in cortical layers (cell counts performed using frames of 400 μm width spanning from L1 to L6 in 9 sections from 3 control mice and 18 sections from 3 *Emx1-cKO* mice; \*\**P* < 0.01, Mann-Whitney test) and in the whole striatal surface (cell counts performed in the whole dorsal striatum in 6 sections from 3 control mice and 7 sections from 3 *Emx1-cKO* mice; *P* = 0.1496, Mann-Whitney test). **d** Representative confocal images showing dendritic spines of GFP-positive L5 neurons from control (*Thy1-GFP-M*) and *Emx1-cKO* (*Thy1-GFP-M; Emx1-cKO*) mice. Scale bar 5 µm. **e** Density of different classes of dendritic spines in control (1688 spines/1135 µm) and *Emx1-cKO* (1308 spines/1220 µm) mice. **f** Coronal brain sections from *GAD67-GFP* control and *Emx1-cKO*-*GAD67-GFP* mice immunostained for TSHZ3. Lower panels are magnifications of the framed areas in the upper images. Scale bars 100 µm. \**P* < 0.02, \*\*\**P* < 0.001 and \*\*\*\**P* < 0.0001, Student’s *t*-test. Data in **b** and **c** are expressed as medians with interquartile range; data in **e** are expressed as means + SEM.

### Cortical excitatory synaptic transmission and corticostriatal synaptic plasticity in *Emx1-cKO* mice

L5 CPNs recorded in slices from *Emx1-cKO* mice show no significant changes in their membrane properties and excitability compared to control (Fig. S4a-e). Action potential (AP)-dependent glutamate release onto L5 CPNs, evaluated by measuring paired-pulse ratio, is also unaffected (Fig. S4f), while both NMDA/AMPA ratio (Fig. S4g) and NMDA-induced currents (Fig. S4h) are significantly reduced, suggesting decreased NMDA receptor-mediated signaling in *Emx1-cKO* mice. The amplitude of AMPA receptor-mediated miniature excitatory postsynaptic currents (mEPSCs) is similar in control and *Emx1-cKO* mice (Fig. S4i), further arguing for the implication of NMDA but not AMPA receptors. Conversely, mEPSC frequency is reduced (Fig. S4i), suggesting decreased AP-independent glutamate release onto L5 CPNs and/or reduced number of active excitatory synapses in *Emx1-cKO* mice, consistent with the decreased spine density on L5 CPNs (Fig. 1d, e).

SSPNs recorded in slices from *Emx1-cKO* mice show electrophysiological properties (Fig. S4A-D) and basal corticostriatal synaptic transmission (Fig. S5e-g) similar to control. However, both long-term potentiation (LTP) and long-term depression (LTD) at corticostriatal synapses are abolished in *Emx1-cKO* mice (Fig. 2), confirming a critical role of *Tshz3* in the functioning of the corticostriatal circuit.

**Figure 2.**
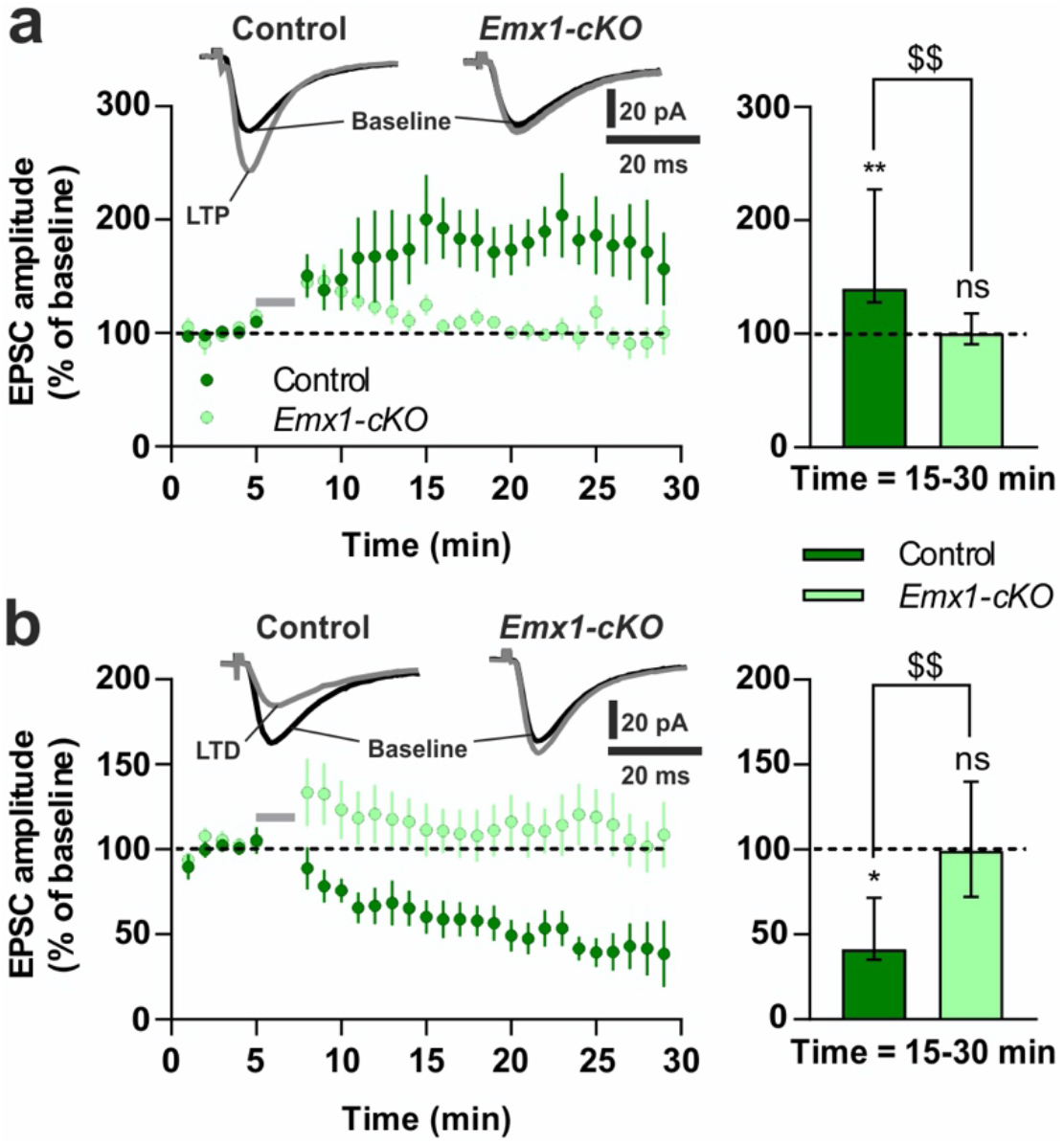
Impaired corticostriatal synaptic plasticity in *Emx1-cKO* mice. LTP **a** and LTD **b** are lost in *Emx1-cKO* mice. Left graphs: time-course (normalized EPSC amplitude expressed as means ± SEM; grey bars represent induction protocols; 2-way ANOVA from 15 to 30 min; LTP: *F*(1,211) = 44.8, *P* < 0.0001; LTD: *F*(1,216) = 153.2, *P* < 0.0001). Traces show EPSCs before (black) and after (grey) LTP and LTD induction protocols. Right graphs: EPSC amplitude at 15-30 min (medians with interquartile range; Wilcoxon matched-pairs signed rank test *vs*. baseline: \**P* < 0.05, \*\**P* < 0.01, ns = non-significant; Mann-Whitney test: ^$$^*P* < 0.01). Data obtained from 17 SSPNs of control and 14 of *Emx1-cKO* mice.

### Conditional deletion of *Tshz3* in cholinergic neurons

Dual immunodetection of CHAT and ß-Gal in *Tshz3*^*+/lacZ*^ mice was performed to analyze the expression of *Tshz3* in brain cholinergic neuron populations. This was preferred to dual immunodetection of CHAT and TSHZ3 since the tissue fixation conditions for obtaining optimal detection of each protein are different, and since TSHZ3 immunodetection provides weaker labeling and higher background than ß-Gal immunodetection. As reported previously ^13^, virtually all SCINs express *Tshz3* (Fig. S6a, h). In contrast, there are no or a little proportion (<30%) of ß-Gal-positive cells within CHAT-positive neurons in the components of the basal forebrain cholinergic system (medial septal nucleus, diagonal band nuclei, *nucleus basalis* of Meynert and *substantia innominata*) (Fig. S6a-d, h). SCINs thus represent the major population of *Tshz3*-expressing cells among the forebrain cholinergic neurons. In addition, there is almost no co-expression of ß-Gal and CHAT in the pedunculopontine (Fig. S6e, f, h) and laterodorsal tegmental nuclei (Fig. S6g, h), which are known to provide cholinergic afferents to several brain areas including the striatum ^19^. Among the other brainstem nuclei, co-expression ranges from poor to extensive, as illustrated in the parabigeminal nucleus and the oculomotor nucleus, respectively (Fig. S6e, f, h).

To address the role of *Tshz3* in cholinergic neurons, *Tshz3*^*flox/flox*^ mice were crossed with *Chat-Cre* mice (*Chat-cKO* model). CHAT is expressed in the brain from early embryonic development and as soon as E18.5 in the striatum ^20^. TSHZ3 immunostaining in *Chat-cKO* mice confirms a significant loss of TSHZ3 in SCINs (Fig. 3a, b), which does not affect the number of striatal CHAT-positive cells (Fig. 3c, d). This result was confirmed using *Chat-Cre;Ai14*^*Flox/+*^ mice (*Chat-Cre*;*Rosa26-STOP-Tomato*) to visualize SCINs in the presence or absence of *Tshz3* (Fig. 3e, f).

**Figure 3.**
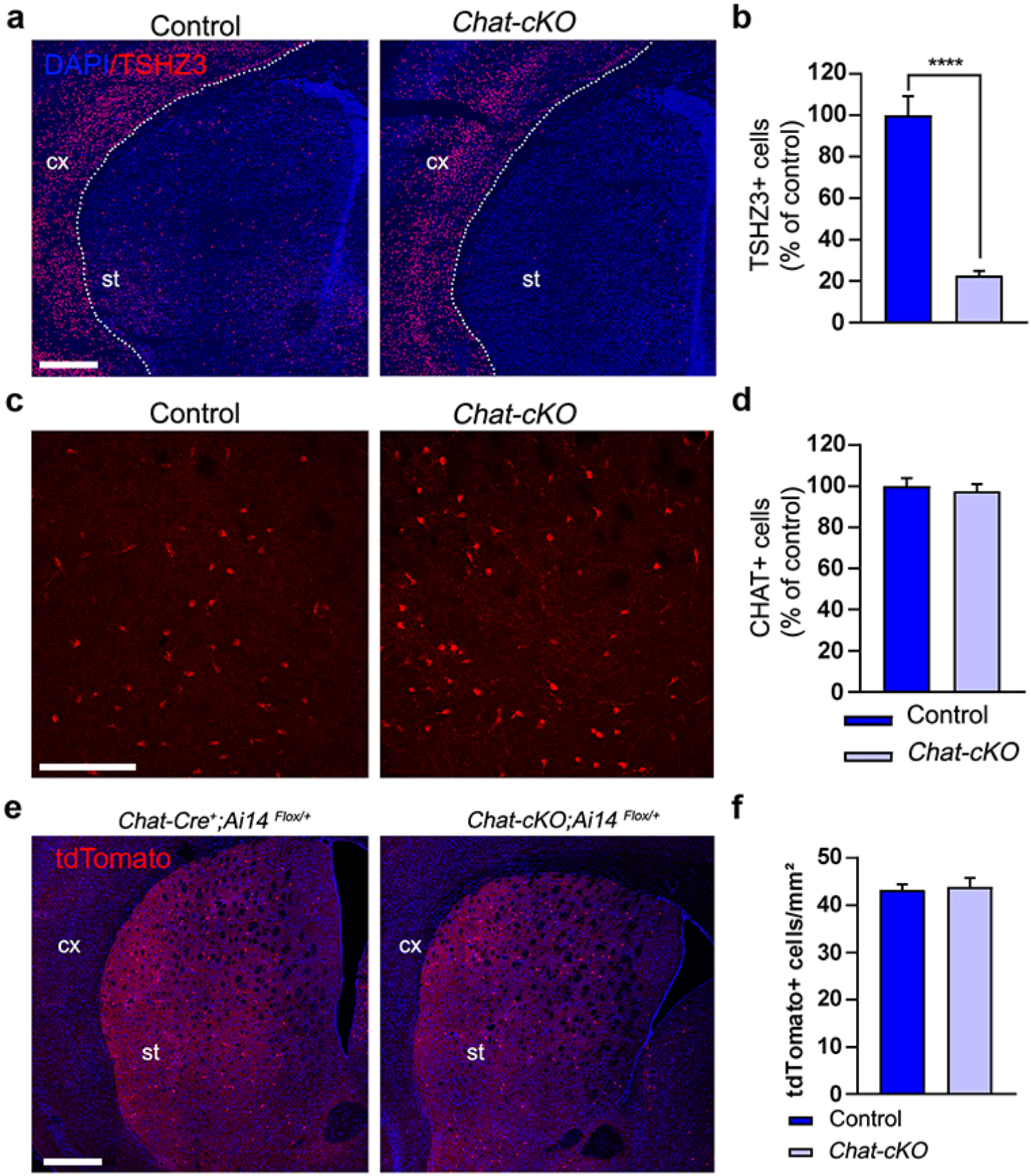
Conditional *Tshz3* deletion in cholinergic neurons. **a** Coronal brain sections from control and *Chat-cKO* mice immunostained for TSHZ3 and counterstained with DAPI. Scale bar 500 µm. **b** Number of TSHZ3-positive cells in the striatum of control and *Chat-cKO* mice; results are expressed as percent of mean control value (15 sections from 3 control mice; 11 sections from 3 *Chat-cKO* mice; \*\*\*\**P* < 0.0001, Student’s *t*-test). **c** Coronal brain sections from control and *Chat-cKO* mice stained for CHAT. Scale bar 200 µm. **d** Number of CHAT-positive SCINs in the striatum of control and *Chat-cKO* mice; results are expressed as percent of mean control value (40 sections from 9 control mice; 53 sections from 11 *Chat-cKO* mice; *P* = 0.6373, Student’s *t*-test). **e** Representative images showing tdTomato fluorescence detection (red) in SCINs of *Chat-Cre;Ai14*^*Flox/+*^ control and *Chat-cKO;Ai14*^*Flox/+*^ mutant mice (coronal sections). cx, cerebral cortex; st, striatum. Nuclei are counterstained with DAPI. Scale bar 500 µm. **f** Density of tdTomato-positive cells in the striatum of *Chat-Cre;Ai14*^*Flox/+*^ control and *Chat-cKO;Ai14*^*Flox/+*^ mutant mice (14 sections from 3 control mice; 12 sections from 3 *Chat-cKO;Ai14*^*Flox/+*^ mice; *P* = 0.6777, Student’s *t*-test). Data in **b, d** and **f** are expressed as means + SEM.

### *Tshz3* loss and SCIN electrophysiological properties

In acute brain slices, SCINs are easily recognizable among the other striatal neurons due to their larger soma ^21^. Moreover, they are the only autonomously active cells, firing action APs with either a regular, irregular or bursting pattern ^22, 23^. SCINs also show a characteristic depolarizing voltage sag in response to the injection of negative current pulses due to the activation of the nonspecific *I*_h_ cation current mediated by HCN channels, which largely contributes to the spontaneous AP discharge characterizing these neurons ^23-25^. To test a possible effect of *Tshz3* loss on these SCIN properties, we measured the mean frequency of spontaneous AP discharge, its regularity [expressed as the coefficient of variation (CV) of the inter-AP intervals], and the amplitude of the sag [expressed as voltage sag ratio (VSR)] in SCINs from *Chat-cKO* mice and control littermates (Fig. 4a-c). We found that SCINs recorded from *Chat-cKO* mice show a significant reduction of both VSR (Fig. 4d) and spontaneous firing frequency (Fig. 4e), as well as an increased CV of inter-AP intervals that suggests a less regular discharge activity (Fig. 4f). The resting membrane potential at steady state is similar between control *vs. Chat-cKO* SCINs (46.64 ± 0.68 *vs*. 45.65 ± 0.64 mV, 56 *vs*. 86 SCINs, respectively; P = 0.305, Student’s *t*-test), while the current-voltage relationship reveals a slight but significant increase of input resistance in *Chat-cKO* SCINs *vs*. control, calculated as the slope of the linear best fit (Fig. 4g; 125.7 ± 4.5 *vs*. 107.5 ± 4.0 MΩ, respectively; *F*(1,911) = 8.816, *P* = 0.0031).

**Figure 4.**
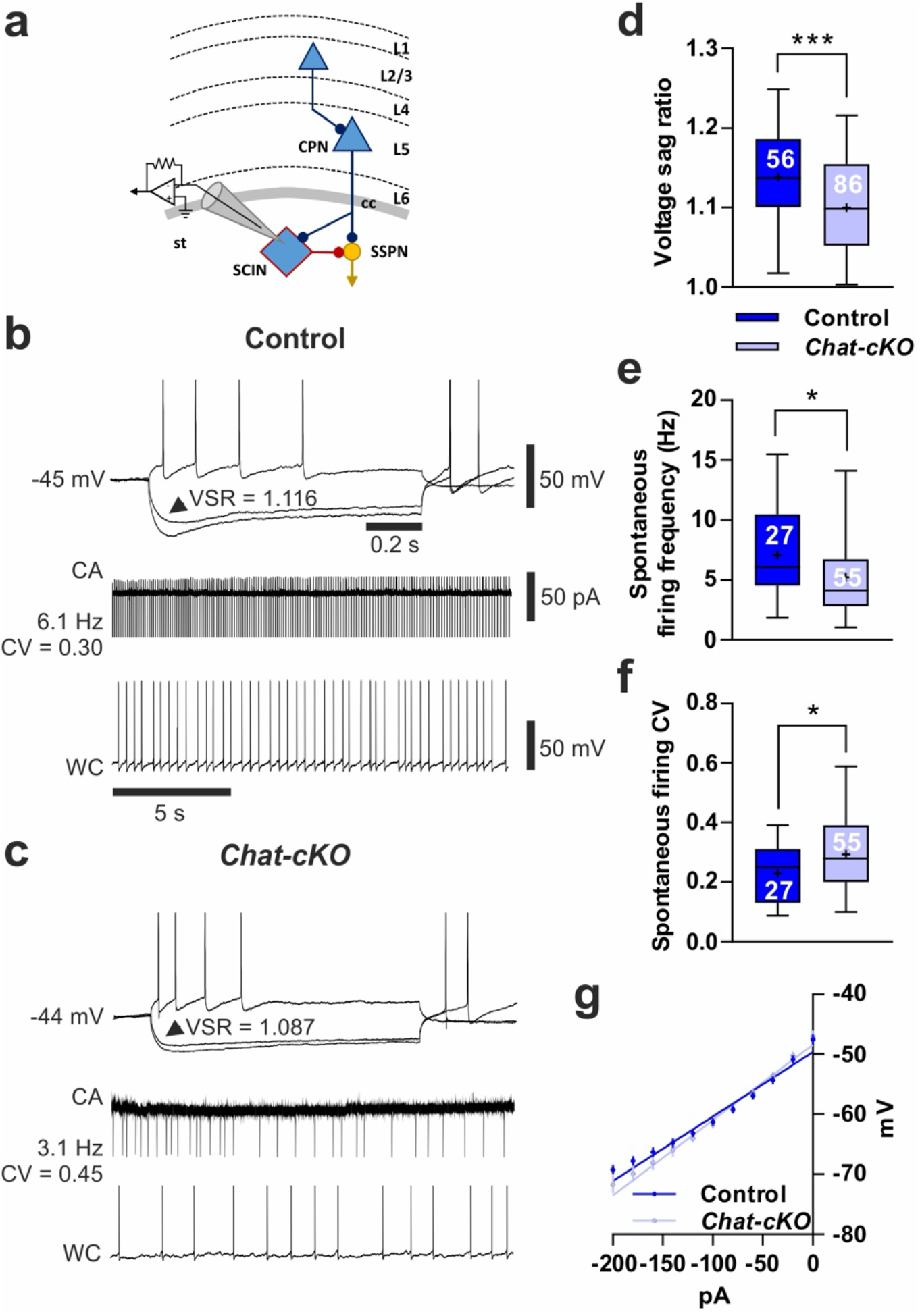
Altered electrophysiological properties of SCINs in *Chat-cKO* mice. **a** Simplified scheme of the corticostriatal circuitry with the recording patch-clamp pipette placed on a SCIN. TSHZ3-expressing neurons are blue (L1-6, cortical layers 1-6; cc, corpus callosum; st, striatum). **b** Sample traces obtained from a representative control SCIN: note the prominent voltage sag in response to -200 and -120 pA hyperpolarizing currents, and the AP firing during a +100 pA depolarizing current (1^st^ line), as well as the sustained and regular firing in cell-attached (CA) and whole-cell (WC) configuration (2^nd^ and 3^rd^ line, respectively). **c** Sample traces obtained from a representative *Chat-cKO* SCIN: compared to **b**, note the smaller voltage sag as well as the less regular, lower frequency spontaneous firing. (b, c) The values of voltage sag ratio (VSR) of the response to -120 pA current injection (arrowhead), as well as the frequency and coefficient of variation (CV) of spontaneous firing of these samples, are reported; spikes have been cut; calibration bars are the same in b and c. Compared to control, SCINs from *Chat-cKO* mice show a significant reduction of mean voltage sag ratio (**d)** and frequency of spontaneous discharge **e**, while the CV of their inter-AP interval is increased (**f)** meaning that their spontaneous firing is more irregular. The number of recorded SCINs in d-f is reported in the graphs. **g** Current-voltage relationship obtained from 51 control and 62 *Chat-cKO* SCINs, and the linear best fit to calculate input resistance (see Results). \**P* < 0.05, \*\*\**P* < 0.001, Student’s *t*-test; data in d-f are expressed as box and whiskers (25th-75th and 5th-95th percentiles, respectively), where bar = median and cross = mean; data in g are expressed as means ± SEM.

### Conditional deletion of *Tshz3* in CPNs or in cholinergic neurons segregates the two core behavioral domains of ASD

For behavioral experiments, only male *Emx1-cKO, Chat-cKO* and control littermate mice were used. Neither *Emx1-cKO* nor *Chat-cKO* mice present visual, auditory and olfactory impairment *vs*. their respective control (Fig. S7). They were then tested for deficits in social behavior, the first core feature of ASD, as well for stereotyped/repetitive patterns of behavior and for restricted field of interests, which are subcategories of the second ASD core feature.

During the habituation phase in the two-chamber test, both *Emx1-cKO* and *Chat-cKO* mice show no significant differences in their exploration of the lured boxes as compared to their respective controls (*P* = 0.14, η^2^ = 0.12, *P* = 0.84, η^2^ = 0.002, respectively Fig. 5a). However, *Emx1-cKO* but not *Chat-cKO* mice show impaired social relationships (Fig. 5). *Emx1-cKO* mice have less preference than their controls for a conspecific (sociability, Fig. 5b) and for an unfamiliar male (social novelty, Fig. 5c), the interaction between genotype and box content being large in each case, as shown by the effect size that exceeds the typical range of variation (Fig. 5d).

**Figure 5.**
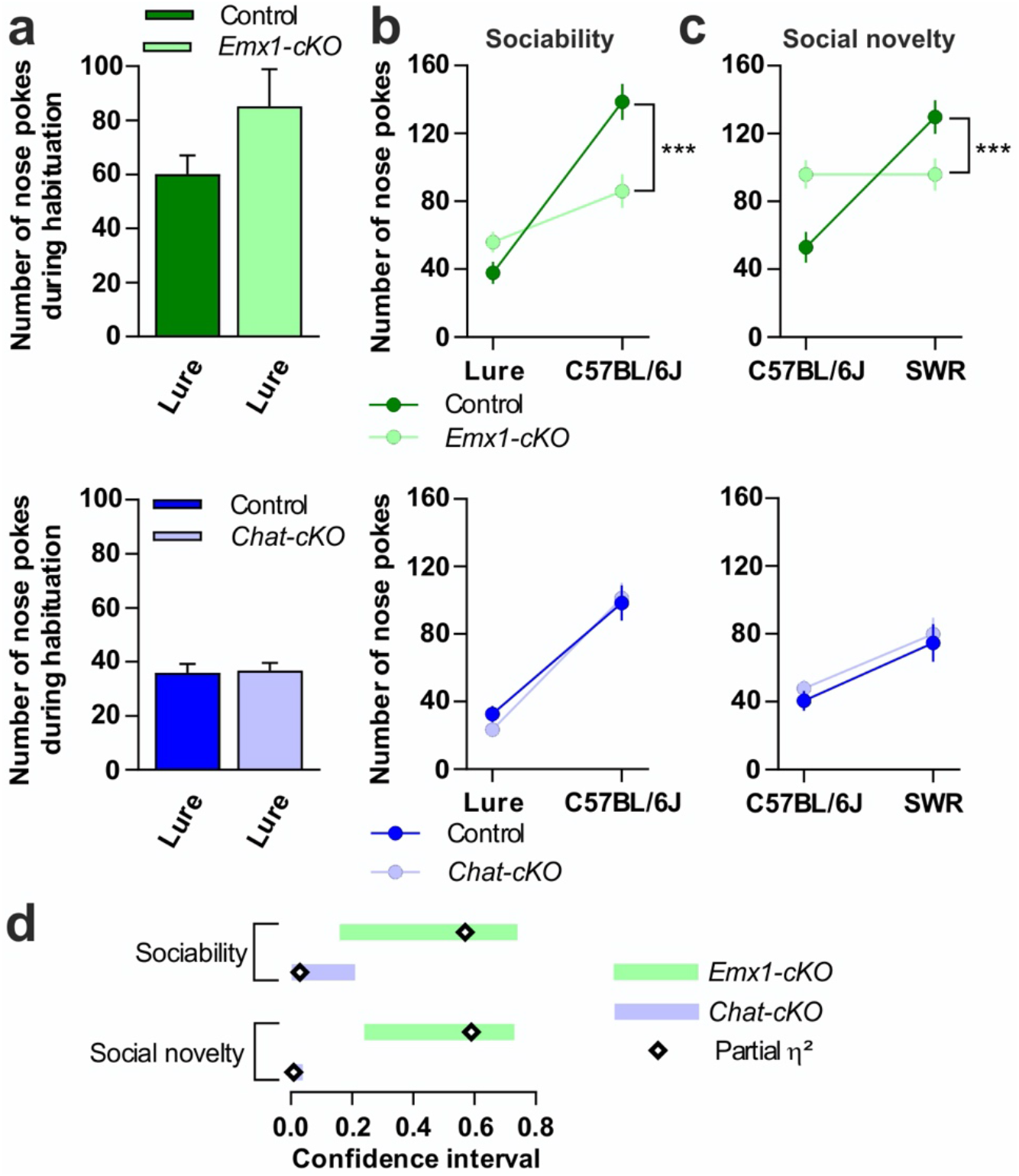
Sociability and social novelty deficits in *Emx1-cKO* but not in *Chat-cKO* mice. **a** Nose pokes during habituation, used as covariate for mixed-design ANCOVAs in **b** and **c. b** Sociability measured as the number of nose pokes against a C57BL/6J male mouse or a lure. *Emx1-cKO* mice (n = 9) *vs*. control (n = 8): *F*_*interaction*_(1,14) = 18.59, *P* < 0.001. *Chat-cKO* mice (n = 12) *vs*. control (n = 9): *F*_*interaction*_(1,18) = 0.55, *P* = 0.47. **c** Interest in social novelty measured as the number of nose pokes against the same C57BL/6J or a SWR mouse. *Emx1-cKO vs*. control: *F*_*interaction*_(1,14) = 19.70, *P* < 0.001. *Chat-cKO vs*. control: *F*_*interaction*_(1,18) = 0.02, *P* = 0.89. **d** Sizes of the difference for *Emx1-cKO* (partial η^2^ = 0.57 and 0.59 for **b** and **c**, respectively) and *Chat-cKO* mice (partial η^2^ = 0.03 and 0.001, respectively) *vs*. their respective control. Data in **a**-**c** are expressed as means ± SEM. \*\*\**P* < 0.001.

Conversely, *Chat-cKO* but not *Emx1-cKO* mice present more stereotyped or repetitive patterns of behavior than their controls, as shown by the marble burying score, time burrowing in a new cage, stereotyped dips on a hole board, and number of leanings in an open field (Fig. 6a-d), with a large effect size (Fig. 6e). Restricted field of interest is impacted neither in *Emx1-cKO* nor in *Chat-cKO* mice (Fig. S8a-c). Finally, hind paw coordination is impaired in *Chat-cKO* but not in *Emx1-cKO* mice (Fig. S8d, e), while spatial learning ability is unaffected in both models (Fig. S8f-i).

**Figure 6.**
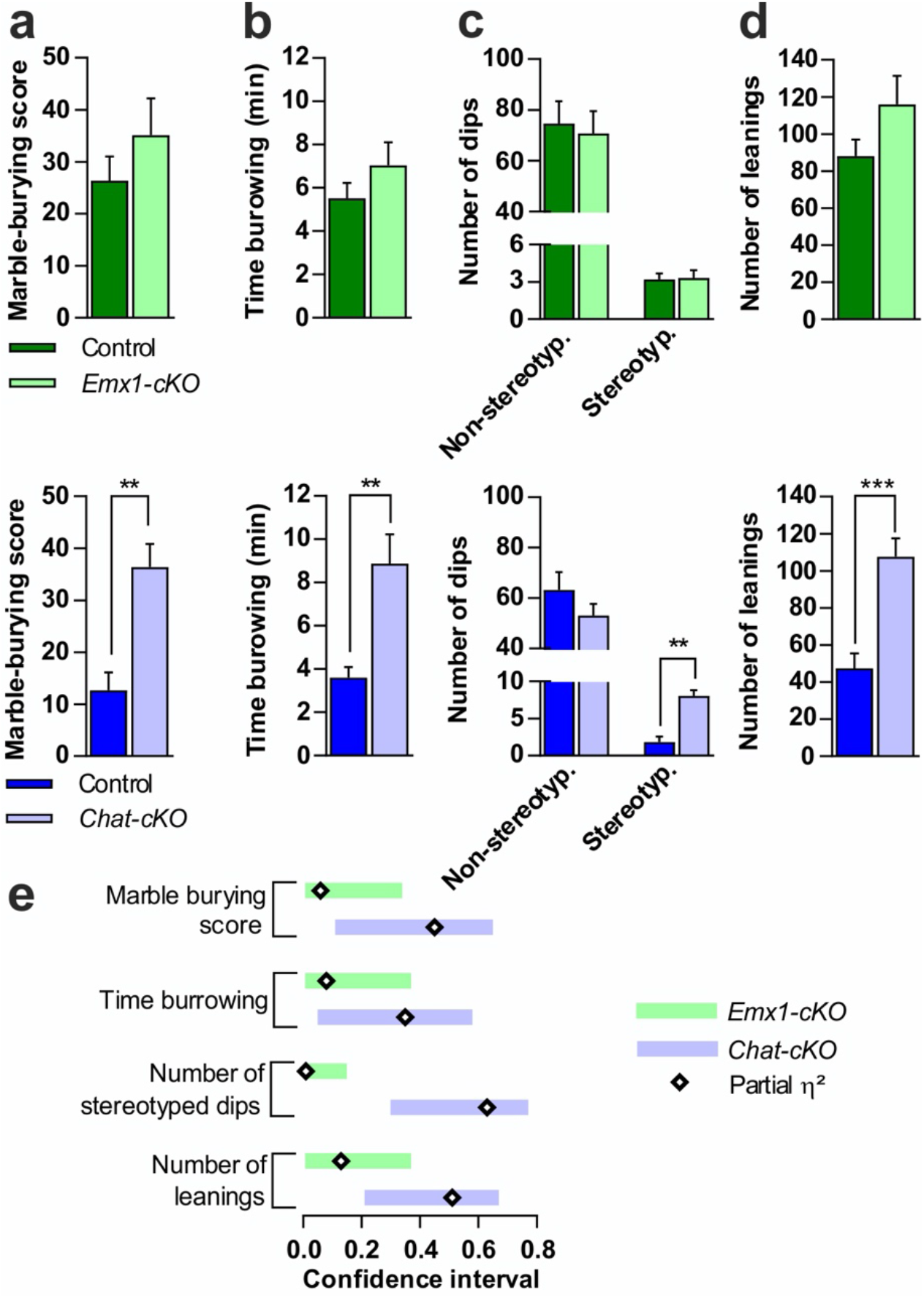
Repeated patterns of behavior in *Chat-cKO* but not in *Emx1-cKO* mice. **a** Marble-burying, *Emx1-cKO*, Student’s-*t*(15) = 1.0, *P* = 0.33; *Chat-cKO, t*(19) = 3.97, *P* = 0.001. **b** Time burrowing, *Emx1-cKO, t*(15) = 1.16, *P* = 0.13); *Chat-cKO, t*(19) = 3.225, *P* = 0.004. **c** Stereotyped dips, *Emx1-cKO, F*_*interaction*_(1,15) = 0.08, *P* = 0.87 (with non-stereotyped dips as covariate, *P* = 0.76); *Chat-cKO, F*_*interaction*_(1,19) = 32.69, *P* = 0.00001 (with non-stereotyped dips as covariate, *P* = 0.24). **d** Number of leanings, *Emx1-cKO, t*(15) = 1.51, *P* = 0.15; *Chat-cKO, t*(18) = 4.35, *P* = 0.0003. **e** Sizes of the difference in *Emx1-cKO* (η^2^ = 0.06, 0.08, 0.13 in a, b and d, respectively, and partial η^2^ = 0.01 in c) and in *Chat-cKO* (η^2^ = 0.45, 0.35, 0.51 in **a, b** and **d**, respectively, and partial η^2^ = 0.63 in c). Sample size of **a, b, c** and d were: 9, 9, 9 and 12 for *Emx1-cKO*; 8, 8, 9 and 11 for their controls; 12, 12, 12 and 11 for *Chat-cKO;* 9, 9, 11 and 8 for their controls. Data in **a**-**d** are expressed as means + SEM. \*\**P* < 0.01 \*\*\**P* < 0.001.

## DISCUSSION

Previous studies showed that haploinsufficiency or postnatal deletion of *Tshz3* results in ASD-relevant behavioral deficits and suggested altered function of the corticostriatal circuitry as a possible substrate ^10, 12^. The present findings point to SCINs as an additional player in the *Tshz3*-linked ASD syndrome. They also provide evidence that targeted conditional deletion of *Tshz3* in either CPNs (*Emx1-cKO*) or cholinergic neurons (*Chat-cKO*) segregates the two core behavioral traits used to diagnose ASD, respectively social behavior deficits and repetitive behavioral patterns, suggesting that alterations in CPNs and in SCINs contribute in a complementary manner to the repertoire of behavioral deficits linked to *Tshz3* deficiency. Restricted field of interest, which defines a sub-category of the second ASD domain, was observed neither in *Emx1-cKO* nor in *Chat-cKO* mice, suggesting that the expression of this deficit in the previously characterized models of *Tshz3* deletion may involve additional players, and/or result from the combined dysfunction of CPNs and SCINs due to the loss of *Tshz3* in both these neuronal types. Learning ability was impacted neither by *Tshz3* postnatal deletion ^12^, nor in *Emx1-cKO* and *Chat-cKO* models.

Among the multiplicity of circuits involved in social behavior, literature points out the crucial role of the cortex ^26, 27^. In particular, corticostriatal and striatal circuit dysfunctions are associated to ASD features, both in patients and in mouse models, with CPNs and SSPNs being highly impacted by mutations of ASD-linked genes ^7, 8, 10, 12, 28, 29^. There is however increasing evidence incriminating interneuron populations of the cortex and the striatum in ASD ^30^. Here, we show that, in the cortex, the ASD-related *Tshz3* gene is expressed not only in CPNs but also in a third of GABA interneurons, while not in cholinergic interneurons. In contrast, in the striatum, the vast majority of *Tshz3*-expressing cells are cholinergic interneurons ^13^. To disentangle the role of CPNs from that of interneurons in the ASD symptoms linked to*Tshz3* deficiency, we generated and characterized *Emx1-cKO* mice. We confirmed the specificity of *Tshz3* deletion in CPNs within the corticostriatal circuit in this model, *Tshz3* expression in cortical and striatal interneurons being maintained. In addition, no change in the numbers and positioning of these interneurons were detected. Interestingly, we found that *Emx1-cKO* mice specifically exhibit impaired social behavior, and that this deficit co-segregates with altered NMDA receptor-mediated transmission in the cortex and disrupted plasticity at corticostriatal synapses. Corticostriatal synaptic plasticity has been deeply characterized, but discrepancies concerning its induction protocols and the underlying molecular and cellular mechanisms ^31^ make it difficult to univocally interpret our results. However, since LTD expression mainly involves presynaptic changes ^32^, its disruption in *Emx1-cKO* mice could be attributable to cortical circuitry defects, such as the observed decrease of NMDA receptor activity in L5 CPNs that could impair corticostriatal output. LTP expression mostly depends upon postsynaptic mechanisms ^32^, but presynaptic NMDA receptors also play a role ^33, 34^. The lack of changes in SSPNs electrophysiological properties or basal corticostriatal transmission rather favors a presynaptic hypothesis to explain this loss of LTP. Moreover, our findings are in line with studies substantiating the involvement of NMDA receptor dysfunction in social deficits associated with ASD in rodent models as well as in patients ^35, 36^. Finally, consistent with the literature linking ASD with changes of dendritic spine density ^37^, we evidence decreased spine density in L5 CPNs of *Emx1-cKO* mice, as in our previous model ^12^. Overall, these data indicate that the loss of *Tshz3* in CPNs induces morphofunctional changes in these neurons and deeply affects corticostriatal plasticity, which might result in altered processing of cortical information and account for the observed social behavior deficits.

We also investigated the contribution of cholinergic neurons in the pathophysiology of *Tshz3*-linked ASD. We show that TSHZ3 is expressed in almost 100% of SCINs, while its expression is absent or partial in the other main brain cholinergic systems. Despite their low number, SCINs have morphofunctional features that place them as key modulators of striatal microcircuits. They play a crucial role in movement control, attentional set-shifting, habit-mediated and goal-directed behavior, and selection of appropriate behavioral responses to changes in environmental contingencies, conferring behavioral flexibility ^38-42^. These interneurons are also involved in basal ganglia-related pathologies such as dystonia, Parkinson’s and Huntington’s disease, Tourette’s syndrome, obsessive compulsive disorder and drug addiction ^43-45^. In contrast, despite the array of data pointing to basal ganglia and cholinergic transmission abnormalities in ASD and in ASD models ^16, 46-50^, to date there is little evidence showing the specific involvement of SCINs: the partial depletion of both SCINs and fast-spiking GABAergic interneurons produces stereotypy and impaired social behavior in male mice ^17^, while total elimination of SCINs results in perseverative behavior that extends to social behavior, rather reminiscent of neuropsychiatric conditions as Tourette’s syndrome or obsessive compulsive disorder ^51^. The present work reveals that targeted *Tshz3* deletion in CHAT-expressing neurons leads to robust stereotyped and repetitive patterns of behavior without impacting social behavior. Given the literature associating drug-induced stereotypies with abnormalities in striatal cholinergic signaling ^52-54^, and the co-expression of CHAT and TSHZ3 in SCINs but not in brainstem cholinergic neurons that are known to project to the striatum ^19^, this behavioral deficit is likely attributable to SCINs. Whereas the number of SCIN in *Chat-cKO* mice is unchanged, suggesting that their generation and viability is not affected, we evidenced modifications in their firing activity and electrophysiological membrane properties. This finding is an addition to the increasing amount of data stressing the complex implication of SCINs in health and diseases ^55^. How the selective loss of *Tshz3* in SCINs leads to these electrophysiological changes, what are their molecular bases and what are the consequences on striatal cholinergic signalling still need to be determined. However, SCINs are important modulators of the two populations of SSPNs forming the “direct” and “indirect” pathways by which the striatum regulates basal ganglia outflow, whose balanced activity is determinant for appropriate action selection ^40, 56^. Thus, the changes in SCIN properties observed here could alter the way they normally respond to salient stimuli and/or reward-associated cues, thereby the way they modulate the transfer of cortical information through the striatum ^38, 39, 57^, as observed after targeted deletion of the transcription factor Er81 in SCINs ^42^. This could underlie the increased stereotyped behaviors observed in *Chat-cKO* mice and, possibly, also in *Tshz3*^*+/lacZ* 10^, as well as in postnatal *Tshz3* cKO ^12^ in which we recently showed that*Tshz3* is lost also in SCINs ^13^. Finally, *Chat-cKO* mice do not show basal exploration deficit, similarly to *Emx1-cKO* mice, but present impaired hind paw coordination, which is in line with motor deficiencies frequently associated with ASD ^58^ and with a study linking partial SCIN ablation with motor incoordination ^59^. Although TSHZ3 is expressed in about 25% of cholinergic neurons of the *nucleus basalis* of Meynert and the *substantia innominata*, the similarity of spatial learning curves of control and *Chat-cKO* mice suggests minor impact of *Tshz3* deletion on the function of the basal forebrain cholinergic system, which is deeply involved in learning and memory processes ^60^.

In conclusion, this study shows that the conditional loss of the ASD-related gene *Tshz3* in CPNs and SCINs does not affect the numbers of these neurons but induces profound changes in their electrophysiological and synaptic properties, associated with specific ASD-like behavioral defects. To our knowledge, it represents the first demonstration in mice models that the two behavioral domains used to diagnose ASD are independent domains that can be underpinned by dysfunction in distinct neuronal subtypes, in this case two components of the corticostriatal circuitry. These findings may open the road to domain-specific pharmacological and behavioral therapies.

## MATERIALS and METHODS

### DATA AVAILABILITY

The data that support the findings of this study are available from the corresponding author upon reasonable request. Raw data (FastQ files) from the sequencing experiment (triplicates from wild-type and *Tshz3*-mutant striatum) and raw abundance measurements for genes (read counts) for each sample are available from Gene Expression Omnibus (GEO) under accession GSE157658, which should be quoted in any manuscript discussing the data.

### MOUSE STRAINS AND GENOTYPING

The *Tshz3*^*lacZ*^, *Tshz3*^*flox/flox*^, *Emx1-Cre, Chat-Cre, Rosa26-STOP-lacZ* and *Ai14* (*Rosa26-STOP-Tomato*), *GAD67-GFP* and *Thy1-GFP* mouse lines have been described previously ^10, 12, 18, 61-66^. Male heterozygous Cre mice were crossed with female *Tshz3*^*flox/flox*^ to generate the two *Tshz3* conditional knockout (cKO) mice models: *Emx1-cKO* and *Chat-cKO* ^18, 64^. Littermate *Emx1-Cre*^*-/-*^ and *Chat-Cre*^*-/-*^ mice were used as respective controls. Animals carrying the *Tshz3*^*flox*^ allele and *Tshz3*^*Δ*^ allele were genotyped as described previously ^12^. Experimental procedures were in agreement with the recommendations of the European Communities Council Directive (2010/63/EU). They have been approved by the “Comité National de Réflexion Ethique sur l’Expérimentation Animale n°14” and the project authorization delivered by the French Ministry of Higher Education, Research and Innovation. (ID numbers 57-07112012, 2019020811238253-V2 #19022 and 2020031615241974-V5 #25232).

### IMMUNOHISTOCHEMISTRY AND HISTOLOGY

All stains were processed on coronal brain sections of postnatal day (P) 28-34 mice. Immunostaining for TSHZ3 alone was performed on cryostat sections of brains immediately removed after anesthesia (ketamine + xylazine, 100 + 10 mg/kg, respectively, i.p.) and frozen in dry ice until use. Before incubation with the antibodies, sections were fixed with 4% paraformaldehyde for 15 min, then washed twice for 5 min in PBS. For TSHZ3 immunostaining and GFP detection, *GAD67-GFP* mice were anesthetized (see above) and transcardially perfused with PBS. Brains were immediately dissected out, post-fixed by immersion 2 hours in 4% paraformaldehyde in PBS, placed in 30% sucrose in PBS overnight and frozen in dry ice until sectioning. For the other stains, mice were anesthetized (see above) and transcardially perfused with 4% paraformaldehyde (PFA) in 0.1 M phosphate buffer. Brains were removed and post-fixed in 4% PFA for at least 2 h before cryostat sectioning (40 µm-thick). Brain sections were washed with PBS and blocked in PBST (0.3% Triton X-100 in PBS) with 5% BSA for 1h at room temperature. Sections were then incubated in primary antibody diluted in blocking solution (PBST, 1% BSA) overnight at 4°C with the following primary antibodies: mouse anti-NeuN (1:500, Millipore, Mab377), rat anti-BCL11B (1:1,000, Abcam, ab18465), goat anti-CHAT (1:100, Millipore, AB144P), rabbit anti-ß-Galactosidase (1:1,000, Cappel, 599762), goat anti-CDP/CUX1 (1:200, Santa Cruz Biotechnology, C20, SC6327) and guinea-pig anti-TSHZ3 (1:2,000; ref. ^61^). Sections were then washed with PBS three times and incubated overnight at 4°C in secondary antibodies diluted 1:1,000 in blocking solution: donkey anti-rabbit Cy3, donkey anti-guinea pig Cy3 and donkey anti-goat Cy3 (Jackson ImmunoResearch Laboratories) and goat anti-mouse Alexa Fluor 488, goat anti-rat Alexa Fluor 555 and donkey anti-goat Alexa Fluor 488 (Life Technologies). Sections were counterstained by 5 min incubation in 300 µM DAPI intermediate solution (1:1,000, Molecular Probes, Cat# B34650). Section were then washed with PBS three times, mounted on Superfrost Plus slides (Fischer Scientific) and coverslipped for imaging on a laser scanning confocal microscope (Zeiss LSM780 with Quasar detection module). Spectral detection bandwidths (nm) were set at 411-473 for DAPI, 498-568 for GFP and 568-638 for Cy3; pinhole was set to 1 Airy unit. Unbiased stereological counting of NeuN, TSHZ3, CUX1, BCL11B, CHAT and ß-Gal positive neurons as well as of GAD-GFP neurons were done from confocal images using ImageJ software (see Figure legends for frame details). Images were assembled using Photoshop 21.2.3.

Cell counts were performed in the dorsal striatum (excluding the nucleus accumbens) and in the surrounding motor and sensorimotor cortex on sections spanning from bregma 0 to +1.18 mm, AP. The whole surface was analyzed for the striatum. For the cortex, counts were performed in frames of 400-μm width either spanning the total thickness of the cortex (NeuN), the thickness of specific layers or divided into 10 bins of equal size for the analysis of the distribution of Gad67GFP-positive cells. For the different cholinergic nuclei, the analyses were performed on sections spanning from bregma +0.62 to +0.38 mm for ms and hdb, -0.34 to -0.8 for si and nbm, +3.8 to -4.16 for 3N, -4.16 to -4.6 for pbg and pptg and -4.72 to - 5.2 for ldtg.

### MORPHOMETRIC AND DENDRITIC SPINE ANALYSIS OF L5 CPNS

We used transgenic mouse lines (P28) expressing *Thy1-GFP* (green fluorescent protein) in L5 CPNs ^66^. *Thy1-GFP-M; Emx1-cKO* were obtained by crossing *Emx1-Cre; Tshz3*^*flox/flox*^ males with *Tshz3*^*flox/flox*^ females heterozygous for *Thy1-GFP*. Analysis of spine density and morphology was performed on stacks from 100 µm-thick vibratome sections (1 µm z-step) on 4 littermate pairs using a Zeiss LSM780 (Oberkochen, Germany) laser scanning confocal microscope (63X objective NA 1.4, 0.03 µm/pixel, voxel size 0.033 µm^2^ x 0.37 µm). Spine counts were obtained from second or third order basal dendritic branches of randomly selected L5 CPNs. Dendrites from 5 to 7 cells were analyzed per animal, providing a cumulated dendrite length > 750 µm for each genotype. Spine identification and density measures were done using NeuronStudio ^67^.

### RT-qPCR

Total RNA from control and *Tshz3* mutant (P28) cerebral cortex was prepared using RNeasy Plus Universal Mini Kit gDNA eliminator (*Qiagen™*) and first strand cDNA was synthesized using iScript Reverse Transcription Supermix kit (Bio-RAD™). Real-time quantitative PCR (RT-qPCR) was performed on a CFX96 qPCR detection system (Bio-RAD™) using *SYBR*® *GreenER*™ qPCR SuperMixes (*Life Technologies*™). RT-qPCR conditions: 40 cycles of 95 °C for 15s and 60 °C for 60 s. Analyses were performed in triplicate. Transcript levels were first normalized to the housekeeping gene *Gapdh*. Primer sequences used for RT-qPCR: *Gapdh* Forward: 5’ GTCTCCTGCGACTTCAACAGCA 3’; *Gapdh* Reverse: 5’ ACCACCCTGTTGCTGTAGCCGT 3’. *Tshz3* Forward: 5’ CACTCCTTCCAGCATCTCTGAG 3’; *Tshz3* Reverse: 5’ TAGCAGGTGCTGAGGATTCCAG 3’.

### ELECTROPHYSIOLOGY

Electrophysiological data were obtained from 57 *Emx1-cKO* and 44 *Emx1-Cre*^*-/-*^ control littermates, and from 16 *Chat-cKO* and 16 *Chat-Cre*^*-/-*^ control littermates, aged P21-28. Procedures were similar to those described previously ^10, 12, 68^. Briefly, acute coronal slices (250 µm-thick) containing cortex and striatum were cut using a S1000 Vibratome (Leica) in ice-cold solution containing (in mM): 110 choline, 2.5 KCl, 1.25 NaH_2_PO_4_, 7 MgCl_2_, 0.5 CaCl_2_, 25 NaHCO_3_, 7 glucose, pH 7.4. Slices were kept at room temperature in oxygenated artificial cerebrospinal fluid (ACSF), whose composition was (in mM): 126 NaCl, 2.5 KCl, 1.2 MgCl_2_, 1.2 NaH_2_PO_4_, 2.4 CaCl_2_, 11 glucose and 25 NaHCO_3_, pH 7.4. Electrophysiological recordings were performed in oxygenated artificial cerebrospinal fluid (ACSF) at 34-35 °C, flowing at ∼2 ml/min. L5 CPNs of the primary motor and somatosensory cortex, and SSPNs and SCINs of the dorsolateral striatum were identified by infrared video microscopy and by their electrophysiological properties ^69, 70^. They were recorded by whole-cell patch-clamp using borosilicate micropipettes (5-6 MΩ) filled with an internal solution containing (in mM): 125 K-gluconate, 10 NaCl, 1 CaCl_2_, 2 MgCl_2_, 0.5 BAPTA, 19 HEPES, 0.3 Na-GTP, and 1 Mg-ATP, pH 7.3 (except for NMDA/AMPA ratio experiments, see below). Electrophysiological data were acquired by an AxoPatch 200B amplifier and pClamp 10.7 software (Molecular Devices, Wokingham, UK). Series and input resistance were continuously monitored by sending 5 mV pulses, and neurons showing ≥ 20% change in these parameters were discarded from the analysis.

### Characterization of CPNs, SSPNs and synaptic transmission

A stimulating bipolar electrode was placed either in the cortex at the level of L4 to activate local fibers and evoke excitatory postsynaptic currents (EPSCs) in L5 CPNs, or in the *corpus callosum* to activate corticostriatal fibers and evoke EPSCs in SSPNs ^12^. Glutamatergic EPSCs were recorded in the presence of 50 µM picrotoxin at a holding potential of -60 mV (CPNs) or -80 mV (SSPNs). Spontaneous miniature EPSCs (mEPSCs) were recorded in the presence of 50 µM picrotoxin and 1 µM tetrodotoxin. Current-voltage (I-V) relationship was obtained in current-clamp mode by injecting hyperpolarizing and depolarizing current steps (ΔI = ±50 pA, 800 ms), and input resistance was calculated by linear regression analysis, i.e. as the slope of the linear best fit of the I-V relationship of each recorded neuron. Rheobase was measured as the minimal injected current (+5 pA increments) capable of eliciting an action potential (AP). For paired-pulse ratio (PPR), EPSC amplitude was measured on 6 averaged traces at each inter-pulse interval. For analyzing mEPSCs, the detection threshold (around 3-4 pA) was set to twice the noise after trace filtering (Boxcar low-pass), and only cells exhibiting stable activity and baseline were considered. For NMDA/AMPA ratio experiments, the internal solution contained (in mM): 140 CsCl, 10 NaCl, 0.1 CaCl_2_, 10 HEPES, 1 EGTA, 2 Mg-ATP and 0.5 Na-GTP, pH 7.3. The AMPA component of the EPSC was measured at the peak at a holding potential of -60 mV, while the NMDA component was measured at +40 mV and 40 ms after the stimulation artifact, when the AMPA component is negligible, as previously described ^12^. Tonic NMDA currents were elicited by bath application of 50 µM NMDA for 60 s, after a stable baseline of at least 120 s; their amplitude was measured by averaging the current values of a 5 s window around the negative peak, compared to baseline; only neurons that were capable of returning to their baseline after washout were considered. EPSC amplitude for monitoring corticostriatal long-term depression and potentiation (LTD and LTP, respectively) was measured on averaged traces (6 per minute) to obtain time-course plots and to compare this parameter before (baseline) and after induction protocols. The induction protocol for corticostriatal LTD consisted of 3 trains at 100 Hz, 3 s duration, 20 s interval, at half intensity compared to baseline ^71^. LTP induction protocol was identical but, during each train, neurons were depolarized to -10 mV to allow strong activation of NMDA receptors ^10, 12, 72^. For a review about corticostriatal LTD and LTP see ^32^.

### Characterization of SCINs

The resting membrane potential (RMP) was measured at the steady state between two consecutive APs. The current-voltage relationship was calculated from the membrane response at the end of current steps from -200 to -20 pA (20 pA steps lasting 800 ms). The voltage sag ratio (VSR) was calculated from the response to a -120 pA current step as the peak voltage drop (sag) against the voltage at the end of the current pulse ^73, 74^. Such relatively small current step was chosen because, with larger steps, the sag amplitude was extremely variable between different SCINs. Spontaneous AP firing was analyzed in terms of discharge frequency (expressed in Hz) and regularity; to quantify this latter parameter, we calculated the coefficient of variation (CV) of the inter-AP intervals. Note that spontaneous AP firing was analyzed only from cell-attached recordings, which were done before switching to whole-cell; in some cases, spontaneous firing was not detectable in cell-attached configuration, thus the number of samples for AP firing analyses is smaller than the whole number of recorded SCINs.

### BEHAVIORAL ANALYSIS

#### Housing conditions

Experiments were conducted blind for the genotypes in P71-87 male *Emx1-cKO* and *Chat-cKO* mice and their respective *Emx1-Cre*^*-/-*^ and *Chat-Cre*^*-/-*^ control littermates. We used males and not female mice because the ambulatory activity of females is impacted by the estrous cycle phases in rodents ^75^ and may bias the results of repetitive behavior measures that are partly dependent on motor activity.

Mice used in studies on social behavior are generally reared in groups of variable size and more rarely in isolation. The choice of our rearing strategy was based on the fact that the measures of social behavior in adult mice depends on the characteristics of the previous interactions that the observed male has experienced with its peers ^3, 76-78^. In the rearing in group strategy, the social behaviors directed towards the tested male can vary according to the genotypes, the androgen levels and the neurotransmitter profiles of the individuals in the groups ^79^. Consequently, the social behavior measured in an individual is the resultant of the individual social ability plus a component corresponding to the interactions of the individual with the other members of the group; this effect varies with the size of the group. In addition, behavioral “contamination” resulting in an impairment of sociability in wild-type mice by cohabitant KO modeling ASD was described ^78^. Such undesirable effect plus the heterogeneity of the measures in mice reared in group should contribute to avoid this strategy for testing social behavior. On the other hand, maintaining the mice socially deprived generates a specific set of agonistic reactions that prevents the measures of social abilities. To circumvent such biases, we have developed an alternative solution for years: each tested male is housed with one female mouse belonging to a single inbred strain ^79^. Here, a cKO or a control male mouse was reared and maintained with CBA/H/Gnc female mice ^3^. Housing was done in transparent 35 × 20 × 18 cm cages with 1-liter poplar woodchip bedding and weekly renewed enrichment (cardboard shelter). The light (07:00-19:00) was 60 lux on the ground of the cages. The temperature was 21.5 ± 0.5°C. Behavioral tests were performed in a dedicated room, the housing cage having been transferred one hour before the beginning of the observations.

#### Assessment of sensory function

Visual, auditory and olfactory integrity is required to ensure the validity of the behavioral data. These sensorial capacities were tested according to previously described protocols ^3, 10, 12, 80^.

*Visual capacities*. The mouse was raised, taken by the tail, and a thin stick was approached to its eyes, without touching the vibrissae. Raising the head was scored 1 and grasping or trying to grasp the pen was scored 2. The test was administered five times and the sum of the scores recorded. Swimming towards a distant shelf in the Morris Water Maze provided an additional assessment of the visual abilities.

*Auditory capacities* were measured using the Preyer’s response. It consists in a pinna twitching and going flat backwards against the head as reaction to sound. It is correlated with the average evoked auditory potential and can be considered as an indicator of auditory acuity ^81-83^. Mice emit vocalizations (less than 20 kHz) and ultrasounds (above 20 kHz) in the presence of a conspecific male. For this reason, we evaluated the responses to stimulations in the ultrasound bandwidth (50 ± 0.008 kHz and 35 ± 0.010 kHz) using commercial dog whistles. The mice received 5 stimulations with each sound. We scored 1 for ear twitching and 2 for a pinna going flat backwards against the head.

*Olfactory ability* to detect an odor was evaluated by an increased time in sniffing a new odor using an olfactory habituation/dishabituation test. Non-social aromas and social odors (urines from C57BL/6J and SWR male mice) were presented individually to each mouse ^84^. The trial was renewed the following day. The individual score was the median time spent.

#### ASD core features

Behaviors modeling the ASD domains as defined by DSM-5 were assessed. The tests were selected based on their strong robustness (reliability from 0.77 to 0.92) and on their high loading scores in a factor analysis ^3^.

*Deficit in social behavior*. A two-chamber test derived from Moy et al., 2004 ^85^ was used to assess sociability and interest in social novelty. The setup and the protocol were detailed previously ^3, 10, 12^. We used a 550 × 550 mm Plexiglas box split in a 150 × 550 mm empty chamber and a 400 × 550 mm chamber containing the two boxes (43 mm diameter, distant from 340 mm) in which the mice or the lure were placed. *Sociability* is operationally defined as the higher number of visits towards the box containing a conspecific versus the one containing a lure (an adult mouse-sized oblong grey pebble), and the *interest in social novelty* as the higher number of visits towards a novel conspecific than towards the familiar one. Loss of social interest and poor interest in social novelty are expected in mice models of ASD. Briefly, the test consisted in a three-period observation, each lasting 10 min: 1) habituation (the two boxes containing lures), 2) sociability (one box containing a lure and the other a C57BL/6J male) and 3) interest in social novelty (one box containing the same C57BL/6J and the other a new SWR male). The behaviors were video-recorded (Viewpoint-Behavior technologies) and the number of nose pokes towards the boxes was counted as measure of the number of visits ^86^.

*Repeated patterns of behavior*. We selected four measures that were highly loaded on the “repetitive patterns of behavior” factor in a factor analysis ^3^: *marble burying* and *time burrowing* in a new cage, *number of stereotyped dips* in a hole-board device, and *number of leanings* in an open field. The protocols used have been previously detailed ^3, 10, 12^.The *marble burying* and *time burrowing* tests quantify perseverating behavior ^87, 88^. *Marble burying* consists in scoring the amount of marbles buried by each mouse in a 30 min session, using a 40 × 18 cm cage with 45 cm-thick litter and containing 20 marbles (9 mm diameter) on the surface of 70 mm-thick dust-free sawdust. Completely buried, 2/3 buried and 1/2 buried marbles were scored 3, 2 and 1, respectively. The *time burrowing* test leans on spontaneous digging and pushing behavior that rodents display when placed into a new home cage. The length of time each mouse spent digging plus pushing was measured. The *number of stereotyped dips* was counted in a hole-board device, consisting in a 40 × 40 cm board with 16 equidistant holes (3.5 cm diameter) each equipped with photo-beams for detecting head dipping. Exploratory head dipping occurs when a rodent is placed on a surface with holes: the mouse puts its head once into one hole of the board. Head dipping is considered stereotyped when the head dips at least twice in the same hole within 2 s. The open field behavior was measured in a circular open field (100 cm diameter and 45 cm high walls) brightly lighted (210 lux on the ground). The ground was virtually divided in three concentric zones of equal surface. The distances walked and the times spent in the open field in the zones were automatically measured via the Viewpoint-Behavior technologies system (http://www.viewpoint.fr/news.php). The observation lasted 20 min. The number of leanings (rearing while leaning) on the walls of the structure was previously validated as a measure of repetitive behavior ^3, 89^. The number of zones crossed is a measure of the narrowness of the field of interest. The total distance walked during the observation period served as covariate for the comparison between cKO mice and their respective controls ^3^.

#### Additional behavioral measures

Motor abnormalities and intellectual disability are not included among the ASD core features while having a noticeable but incomplete prevalence in ASD patients (≤79% and ∼45%, respectively ^58^). In this connection, two additional tests were conducted.

##### Hind paw coordination

A mouse was first trained to cross a smooth bar (50 × 5 × 5 cm) with large platforms on each extremity. The trained mouse was then placed on the central platform (3 × 5 cm) of a notched bar (100 cm) formed of 1.5 cm deep carvings regularly spaced (2 cm). The task consisted in ten bar crossings from the central to an extremity platform. The experimenters on each side of the setup counted the left and right hind paw slips according to ^90^.

##### Spatial learning

The Morris water maze provides measures of the ability of rodents to solve spatial learning problems, namely the ability to find a submerged resting platform concealed beneath opaque water. The platform is a glass cylinder (66 mm diameter, 9 mm beneath the surface of the water) positioned 23 cm from the edge of a 100 cm diameter circular tank filled with water at 26 ± 1°C and the light at 70 lux on the surface. Each mouse performed 7 blocks of 4 trials each: one block on day 1, and two blocks daily (one in the morning and one in the afternoon) for 3 successive days. A trial was stopped after 90 s if the mouse failed to reach the platform. We considered that the mouse had reached the platform when it stayed on the platform for 5 s at least. We presented a small metal shelf to the mouse 5 cm above the platform at the end of each trial of the first block (shaping). The mouse climbed on it and was transferred in a cage with dry sawdust for 120 s. We had previously assigned 4 virtual cardinal points to the tank, each being the starting point for a trial. The starting point for each trial was chosen randomly and within a block the mouse never started more than once from the same virtual cardinal point. We measured 1) the time to reach the hidden platform and 2) the cumulative distance to the center of the platform during swimming. The second measure eliminates possible bias resulting from floating during the trial. The time to reach the platform and the distance were automatically measured by a video tracking setup (Viewpoint-Behavior technologies), each over the 7 blocks. Strains can achieve different performance levels between blocks, but without a cumulative reduction in the time to reach the platform, which is the criterion to identify learning process. We computed the slopes of the learning curves, a negative slope indicating learning behavior ^91^. The strategy was used for both the time to reach the hidden platform and the cumulative distance to the center of the platform. The probe-test procedure, conducted after removing the platform, was done 24 h after block 7 to meet the requirements for reference memory ^92^ and lasted 90 s. The mouse was placed in the center of the tank, and we measured the time of first crossing the virtual annulus corresponding to the location of the platform. To check whether the differences in the time to reach the platform were due to vision and/or swimming abilities rather than learning ability, we also tested groups of naïve *Emx1-cKO* and *ChAT-cKO* mice, and their respective control, to the visible platform version of the test, in which the platform is 5 mm above non-opacified water.

### STATISTICS

#### Immunohistochemistry

Data were analyzed by Prism 7.05 (GraphPad Software, USA). Sample sizes, tests used, and *P* values are reported in Figure legends. The significance threshold was set at *P* < 0.05.

#### RT-qPCR

Statistical analysis for was performed by unpaired Student’s *t*-tests using the qbasePLUS software version 2 (Biogazelle). The significance threshold was set at *P* < 0.05.

#### Electrophysiology

Statistical analysis was performed by Prism 7.05 (GraphPad Software, USA). Student’s *t*-test or two-tailed Mann-Whitney test was used for comparing two data sets when passing or not D’Agostino & Pearson’s normality test, respectively. Two-way ANOVA was used to analyze the influence of 2 categorical variables. 2-samples Kolmogorov-Smirnov test was used to compare cumulative distributions. Sample sizes (n) reported in Figure legends refer to the number of recorded neurons. The significance threshold was set at *P* < 0.05. Tests used, *P* values and sample sizes are indicated in the Figure.

#### Behavior

Data were processed by *Statistical Package for the Social Sciences* [SPSS software, version 25 ^93^]. The same statistical designs were used to compare *Emx1-cKO* and *ChAT-cKO* mice to their respective controls. Non-parametric statistics were chosen when the assumption of normality was rejected.

#### Impairment of social behavior

To analyze data from each social phase of the two-chamber test (sociability and interest for social novelty), a mixed design analysis of covariance (ANCOVA) was used including the genotype as fixed factor, the box content as repeated measure, with measure of activity during habituation as covariate. A significant interaction between genotype and box content indicates that social behavior differs between the cKO and its control group.

#### Repetitive patterns of behavior and motor performance

The difference between two independent groups (cKO and its control group) was tested by an unpaired two-sample Student’s *t*-test in each case where it was not necessary to include a covariate in the statistical design (i.e., stereotyped behavior: marble-burring score, time burrowing, number of leanings; motor behavior: number of hind paw slips). For measures of stereotyped dips, on which the activity level could have an impact, an analysis of covariance (ANCOVA) was performed, using the genotype as fixed factor (cKO *vs*. respective control) and non-stereotyped dips as covariate.

#### Sensorial abilities

Comparison of the visual and auditory capacities of the cKO and their respective controls were conducted using a Student’s *t*-test. Mixed repeated measures ANOVA, with genotype as fixed factor and 15 odors as repeated measures, was used to compare cKO and their respective controls for olfactory capacities.

#### Spatial learning

The statistical design was the same for the time to reach the platform and the cumulative distance to the center of the platform in the Morris water maze test. Differences between the 7 blocks were tested either with Friedman’s ANOVA, a non-parametric version of one-way repeated measures ANOVA, or with two-way repeated measures mixed ANOVA design, with blocks as repeated-measures variable and cKO *vs*. control as between-group variable. Learning may be deduced from within-bloc statistical difference and reduced time to reach the platform from one bloc to the next. The slope of the median values of the four trials in each of the seven blocks was calculated for each mouse. The median slopes for the cKO and their respective controls, as well as the time to reach the virtual platform (probe test) and the visible platform, were compared with a Student’s *t*-test

#### Effect size

Effect sizes are expressed as *η*^*2*^ or as partial *η*^*2*^ with 95% confidence interval ^93, 94^

## Ethic Statement

The animal study was reviewed and approved by the “*Comité National de Réflexion Ethique sur l’Expérimentation Animale n°14*” and the project authorization delivered by the French Ministry of Higher Education, Research and Innovation. (ID numbers 57-07112012, 2019020811238253-V2 #19022 and 2020031615241974-V5 #25232) and were in agreement with the European Communities Council Directive (2010/63/EU).

## Acknowledgements

Behavioral testing was performed at the mouse facility of the Marseille Medical Genetics (MMG) UMR1251 Aix Marseille Univ, INSERM. Microscopy was performed at the imaging platform of the IBDM and we acknowledge France-BioImaging/PiCsL infrastructure (ANR-10-INSB-04-01). This work has received support from the French government under the Programme “Investissements d’Avenir”, Initiative d’Excellence d’Aix-Marseille Université via A*Midex funding (NeuroMarseille Institute, AMX-19-IET-004; MarMaRa Institute, AMX-19-IET-007), and ANR (ANR-17-EURE-0029). We wish to thank the IBDM mouse facility.

## Author Contribution

X.C., J.M. and P.S. performed the histological experiments and the quantitative analyses; M.C. and P.L.R. conducted the behavioral experiments and analyzed the resulting data; Y.B., L.B., J.M. and D.C. performed patch-clamp experiments and P.G. analyzed electrophysiological data; M.M. performed dendritic spine imaging and counting; A.F. performed RT-qPCR; X.C. and J.M. generated and maintained transgenic mouse lines; X.C., L.F., P.G. and L.K.-L.G. conceived the project, supervised the work and wrote the paper with the contribution of M.C. and P.L.R; all authors read and approved the final manuscript.

## Supplementary Figures

**Figure S1.**
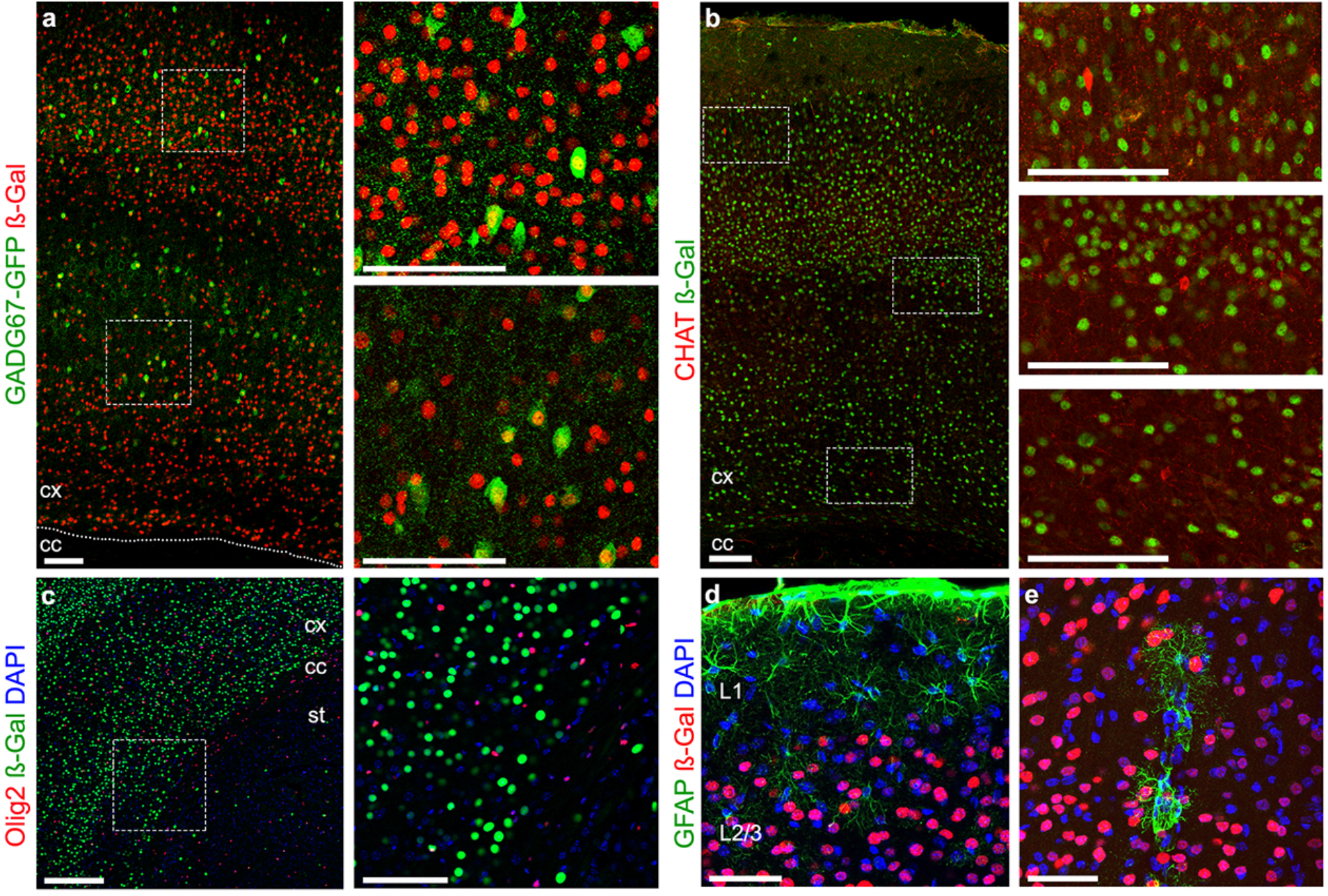
TSHZ3 expression in interneurons and glial cells in the cerebral cortex. (**a**-**e**) Coronal brain sections. **a** *Tshz3* expression as β-Gal staining in *Tshz3*^*+/lacZ*^; *GAD67-GFP* mouse brain. The two images on the right are magnifications of the framed areas in A. Scale bars 100 µm. **b** Double immufluorescence staining for β-Gal and CHAT. The framed areas in (**b**) are magnified on the right. Scale bars 100 µm. **c** Double immufluorescence staining for Olig2 and ß-Gal (left) and detail of the framed area (right). Scale bars 100 µm. (**d, e**) Double immufluorescence staining for GFAP and ß-Gal. Scale bars 100 µm (**d)** and 50 µm (**e)**. Nuclei in **c**-**e** are counterstained with DAPI. cc, corpus callosum; cx, cerebral cortex; st, striatum.

**Figure S2.**
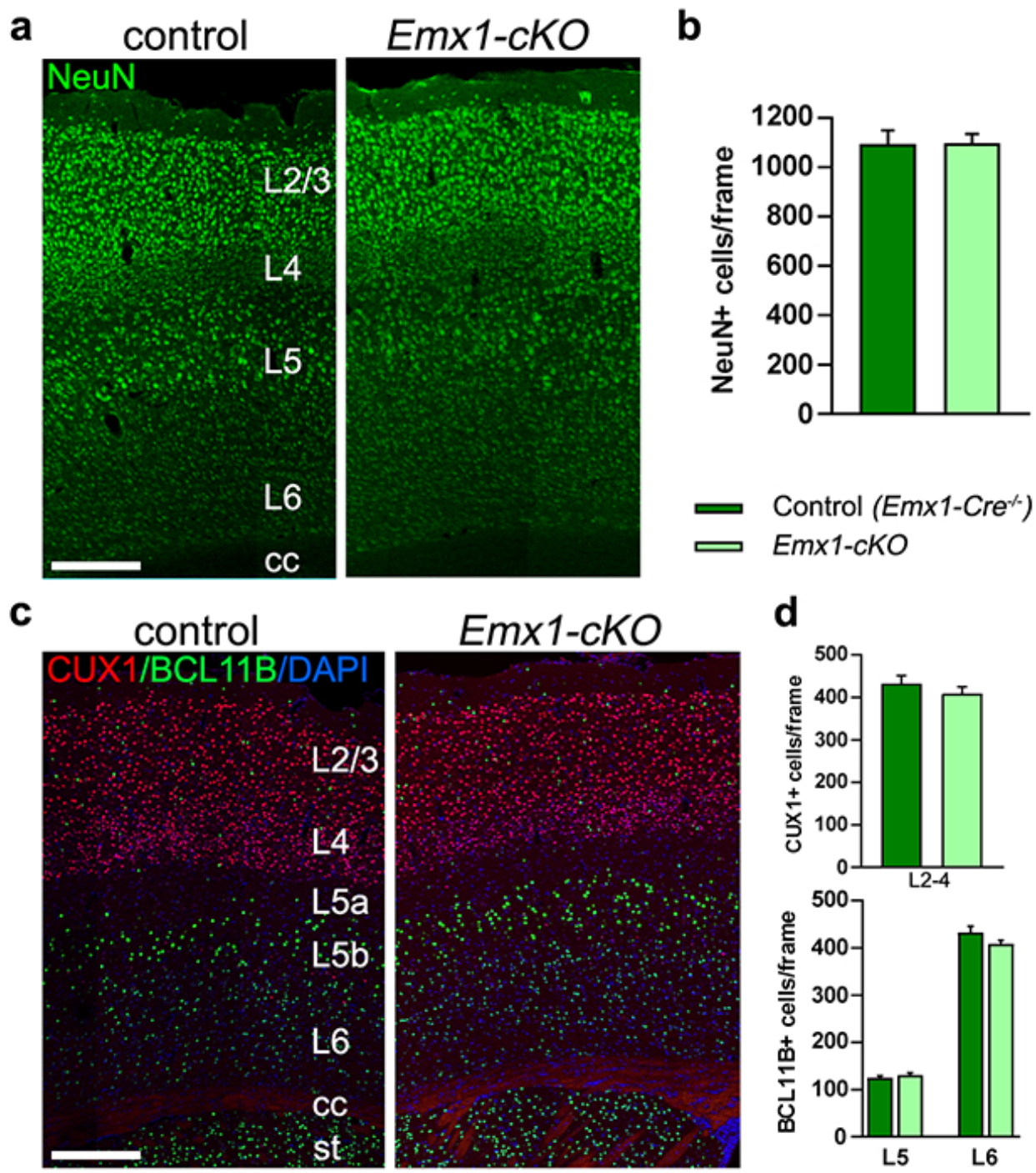
Cortical layering is preserved in *Emx1-cKO* mouse brain. **a** Coronal brain sections from *Emx1-cKO* and control mice immunostained for NeuN detection. Scale bar 250 µm. **b** Number of NeuN-positive cells counted in frames of 400 μm width spanning the entire cortical thickness of control and *Emx1-cKO* mice. No genotype difference is found (11 sections from 3 mice per genotype; *P* = 0.9488, Student’s *t*-test). **c** Coronal brain sections from *Emx1-cKO* and control mice immunostained for CUX1 and BCL11B. Nuclei are counterstained with DAPI. Scale bar 100 µm; cc, corpus callosum; st, striatum; L, layer. **d** Number of CUX1-positive cells in L2-4 and of BCL11B-positive cells in L5 and L6 in control and *Emx1-cKO* mice. No genotype difference is found (BCL11B-positive cells: 14 sections from 3 control mice and 18 sections from 3 *Emx1-cKO* mice; CUX1-positive cells: 28 sections from 4 control mice and 21 sections from 4 *Emx1-cKO* mice; countings were performed in cortical frames of 400 μm width; *P* = 0.3207 (L2/3), *P* = 0.4007 (L5) and *P* = 0.1180 (L6), Student’s *t*-test). Data are expressed as means + SEM.

**Figure S3.**
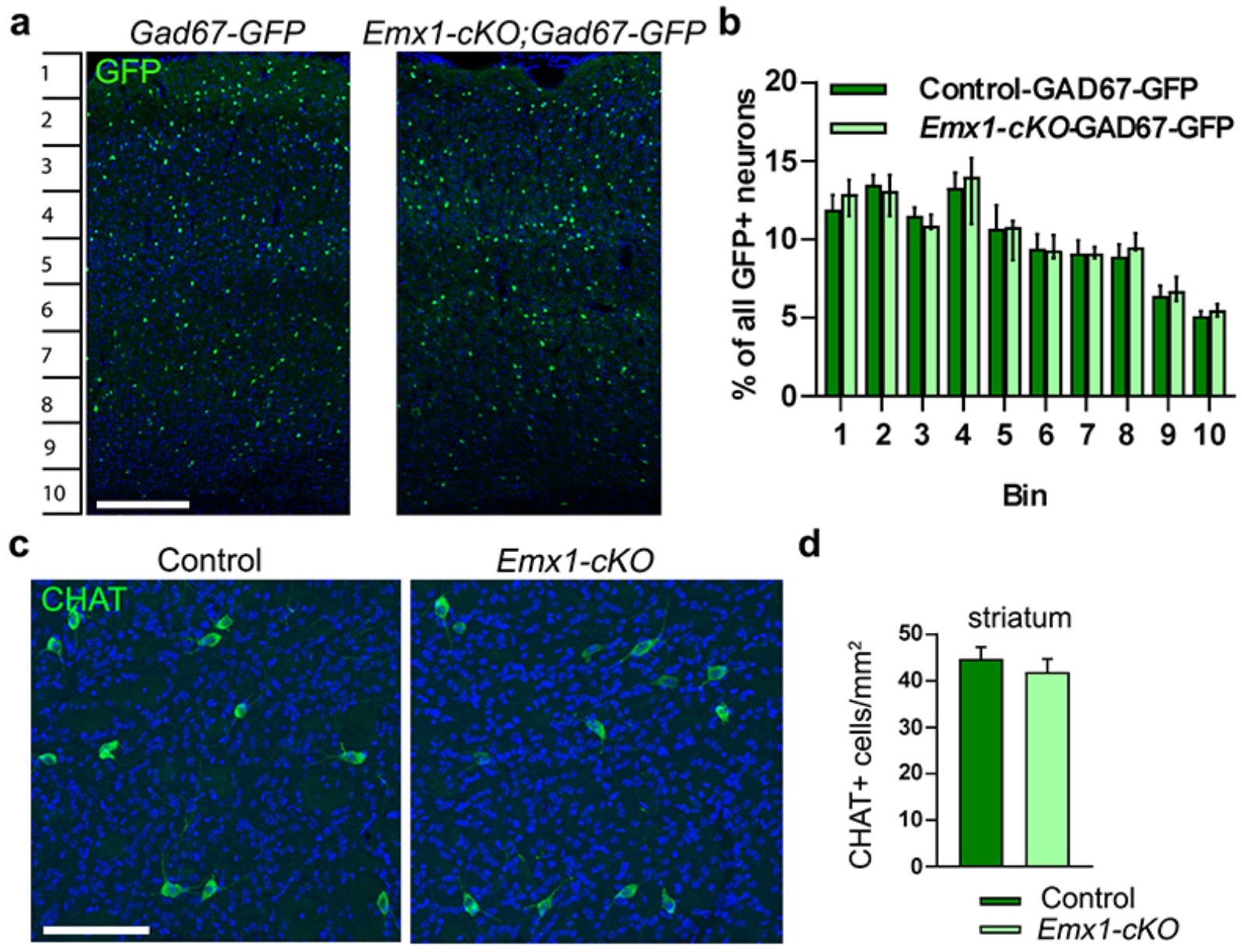
Loss of *Tshz3* in *Emx1-cKO* mice does not affect the numbers of cortical GABAergic and striatal cholinergic interneurons. Representative images **a** and quantitative analysis **b** showing the distribution of GAD67-GFP-positive cells in the cerebral cortex in coronal brain sections from *GAD67-GFP* control and *Emx1-cKO-GAD67-GFP* mice. Scale bar in A 250 µm. Data in b are expressed as percent of total GFP-positive cells per bin (37 sections from 5 control mice; 41 sections from 7 *Emx1-cKO* mice; *F*_*genotype*_(1,100) = 0.00006, *P* = 0.994, *F*_*interaction*_(9,100) = 0.381, *P* = 0.942, 2-way ANOVA). Images of CHAT immunostaining **c** and analysis of the density of CHAT-positive cells **d** in coronal brain sections at striatal level of control and *Emx1-cKO* mice. Scale bar 100 µm (18 sections from 3 control and 3 *Emx1-cKO* mice, respectively; *P* = 0.465, Student’s *t*-test). Data in **b** are expressed as median with interquartile range; data in **d** as means + SEM.

**Figure S4.**
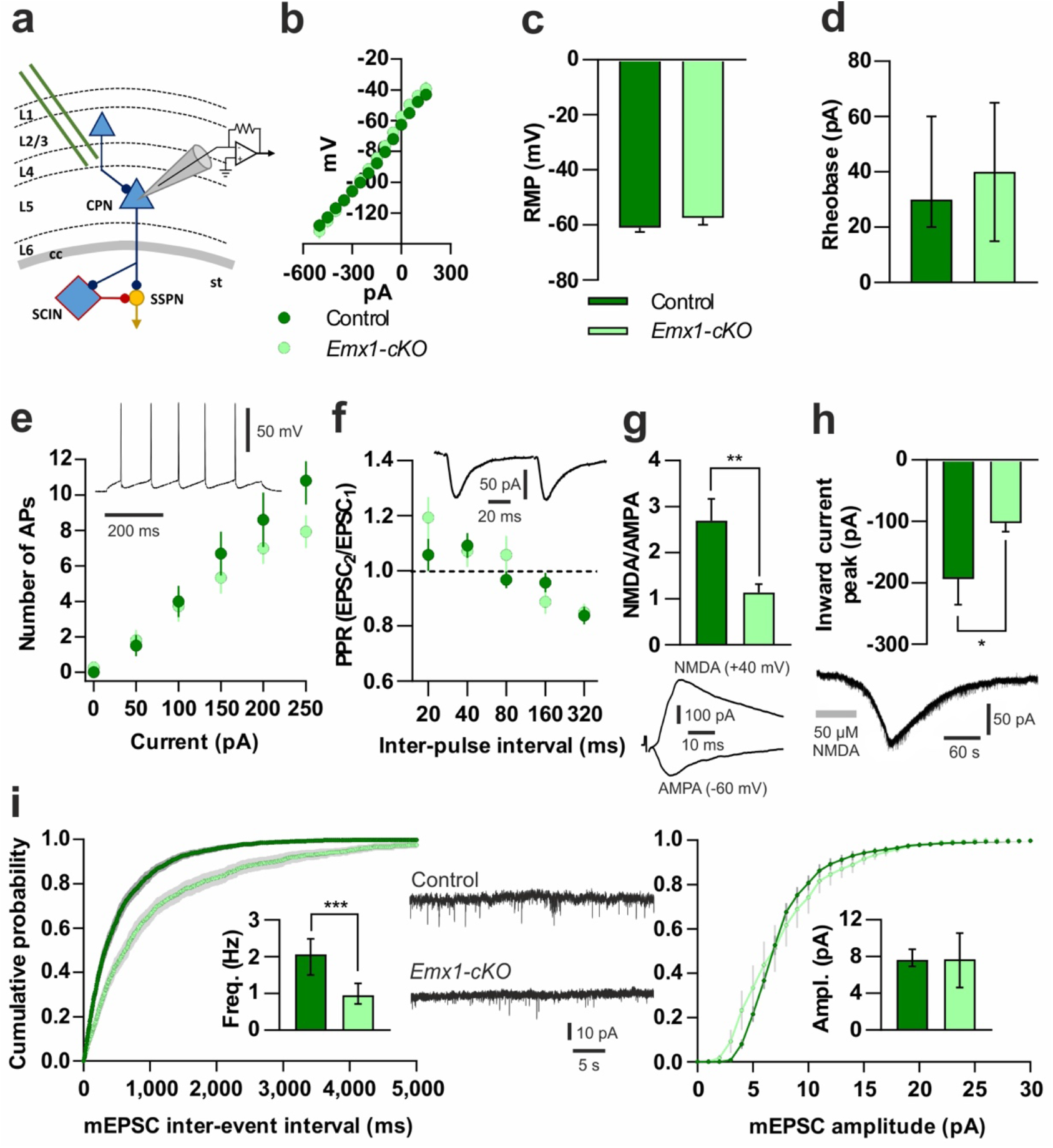
Electrophysiological characterization of L5 CPNs and basal cortical synaptic transmission. **a** Simplified scheme of the corticostriatal circuitry with the recording patch-clamp pipette placed on a L5 CPN and the stimulating electrode placed in L4. TSHZ3-expressing neurons are blue (L1-6, cortical layers 1-6; cc, corpus callosum; st, striatum). **b** Current-voltage relationship recorded from CPNs of *Emx1-cKO* mice and littermate controls show similar slopes and input resistance (148.9 ± 13.3 *vs*. 151.3 ± 11.6 MΩ, respectively; n = 21 and n = 28, respectively; *P* > 0.05, Student’s *t*-test). **c** Resting membrane potential (RMP; n = 28-38) and **d** rheobase (n = 11-21) do not significantly differ between control and *Emx1-cKO* CPNs (*P* > 0.05 for both; Student’s *t*-test and Mann-Whitney test, respectively). **e** The number of action potentials (APs) emitted by control (n = 10) and *Emx1-cKO* (n = 15) CPNs in response to increasing current injections is similar (2-way ANOVA: genotype *F*(1,138) = 3.068, *P* = 0.0821; interaction *F*(5,138) = 0.9349, *P* = 0.4605; multiple *t*-tests: *P* > 0.05). The trace shows an example of AP firing during a 100 pA, 500 ms current step. **f** Paired-pulse ratio (PPR) is not significantly different between control (n = 19) and *Emx1-cKO* (n = 14) CPNs (2-way ANOVA: genotype *F*(1,155) = 0.901, *P* = 0.344; interaction *F*(4,155) = 1.431, *P* = 0.2263). The trace shows an example of paired EPSCs (80 ms inter-pulse interval). **g** NMDA/AMPA ratio is significantly decreased in CPNs of *Emx1-cKO* mice compared to control (n = 15 for each genotype, \*\**P* < 0.01, Student’s *t*-test). Traces show an example of a NMDA- and an AMPA receptor-mediated EPSC recorded from the same CPN at +40 and -60 mV, respectively. **h** The tonic inward currents induced by bath application of NMDA (50 µM, 60 s) are significantly smaller in CPNs from *Emx1-cKO* mice compared to control (n = 15 and n = 14, respectively; \**P* < 0.05, Student’s *t*-test). The trace shows a sample response of a CPN (sEPSCs have been cut) to NMDA bath application (grey bar). **i** The distribution of mEPSC inter-event intervals is significantly different between control (n = 12) and *Emx1-cKO* (n = 11) CPNs (*P* < 0.0001, 2-samples Kolmogorov-Smirnov test), as well as their median frequency (inset) (\*\*\**P* < 0.001, Mann-Whitney test). Conversely, both the distribution and the median values of mEPSC amplitude are similar in control and *Emx1-cKO* CPNs (*P* > 0.05, 2-samples Kolmogorov-Smirnov test and Mann-Whitney test). Cumulative plots represent mean values (light and dark green) and SEM (grey). Traces show sample mEPSCs recorded from control and *Emx1-cKO* CPNs. Data in **b, c, e**-**h** and in **i** (cumulative plots) are expressed as means ± SEM; data in **d** and in **i** (insets) are expressed as medians with interquartile range.

**Figure S5.**
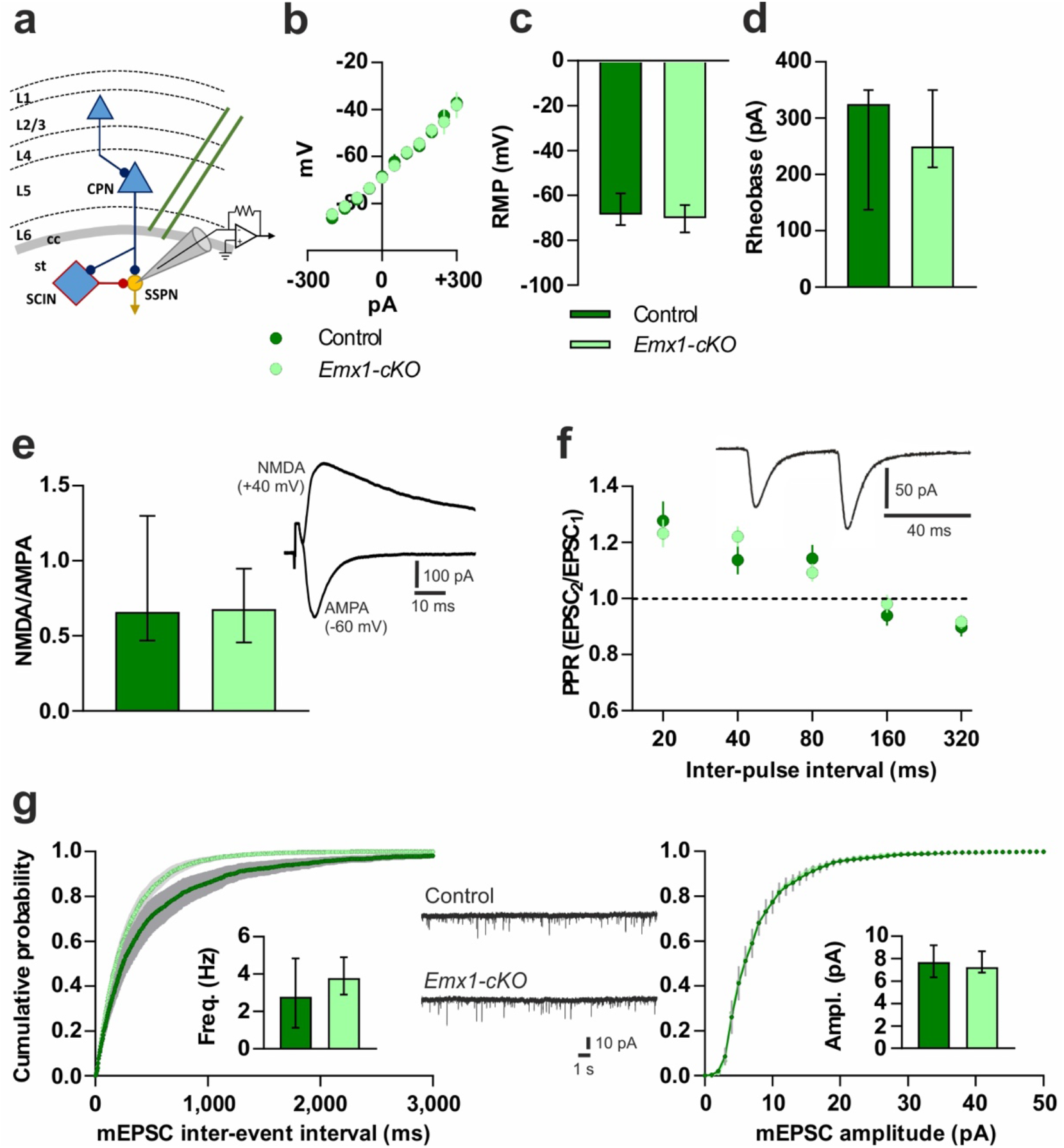
Electrophysiological characterization of SSPNs and basal corticostriatal synaptic transmission. **a** Simplified scheme of the corticostriatal circuitry with the recording patch-clamp pipette placed on a SSPN and the stimulating electrode placed on the cc. TSHZ3-expressing neurons are blue (L1-6, cortical layers 1-6; cc, corpus callosum; st, striatum). **b** Current-voltage relationship recorded from SSPNs of control and *Emx1-cKO* mice provide similar slopes and input resistance (97.4 ± 2.3 *vs*. 93.0 ± 2.1 MΩ, respectively; n = 7 and n = 15, respectively; *P* = 0.1862, Mann-Whitney test). **c** Resting membrane potential (RMP) and **d** rheobase are not significantly different between control (n = 7) and *Emx1-cKO* (n = 15) SSPNs (*P* > 0.05, Mann-Whitney test). **e** NMDA/AMPA ratio is similar between control (n = 11) and *Emx1-cKO* (n = 12) SSPNs (*P* > 0.05, Mann-Whitney test); traces in **e** show an example of an NMDA receptor- and an AMPA receptor-mediated EPSC recorded from the same SSPN at +40 and -60 mV, respectively. **f** Paired-pulse ratio (PPR) is similar between control (n = 18) and *Emx1-cKO* (n = 24) SSPNs (2-way ANOVA: genotype *F*(1,162) = 0.1135, *P* = 0.7367; interaction *F*(4,162) = 0.8429, *P* = 0.4999). The trace shows an example of paired EPSCs (40 ms inter-pulse interval). **g** The distribution of mEPSC inter-event intervals is significantly different between control (n = 8) and *Emx1-cKO* (n = 7) SSPNs (*P* < 0.001, 2-samples Kolmogorov-Smirnov test), but their median frequency (inset) is similar (*P* > 0.05, Mann-Whitney test). Both the distribution and the median value of mEPSC amplitude are not significantly different between control and *Emx1-cKO* SSPNs (*P* > 0.05, 2-samples Kolmogorov-Smirnov test and Mann-Whitney test). Cumulative plots represent average values (light and dark green) and SEM (grey). Traces show sample mEPSCs recorded from control and *Emx1-cKO* SSPNs. Data in **b, f** and **g** (cumulative plots) are expressed as means ± SEM; data in **c**-**e** and **g** insets are expressed as medians with interquartile range.

**Fig. S6.**
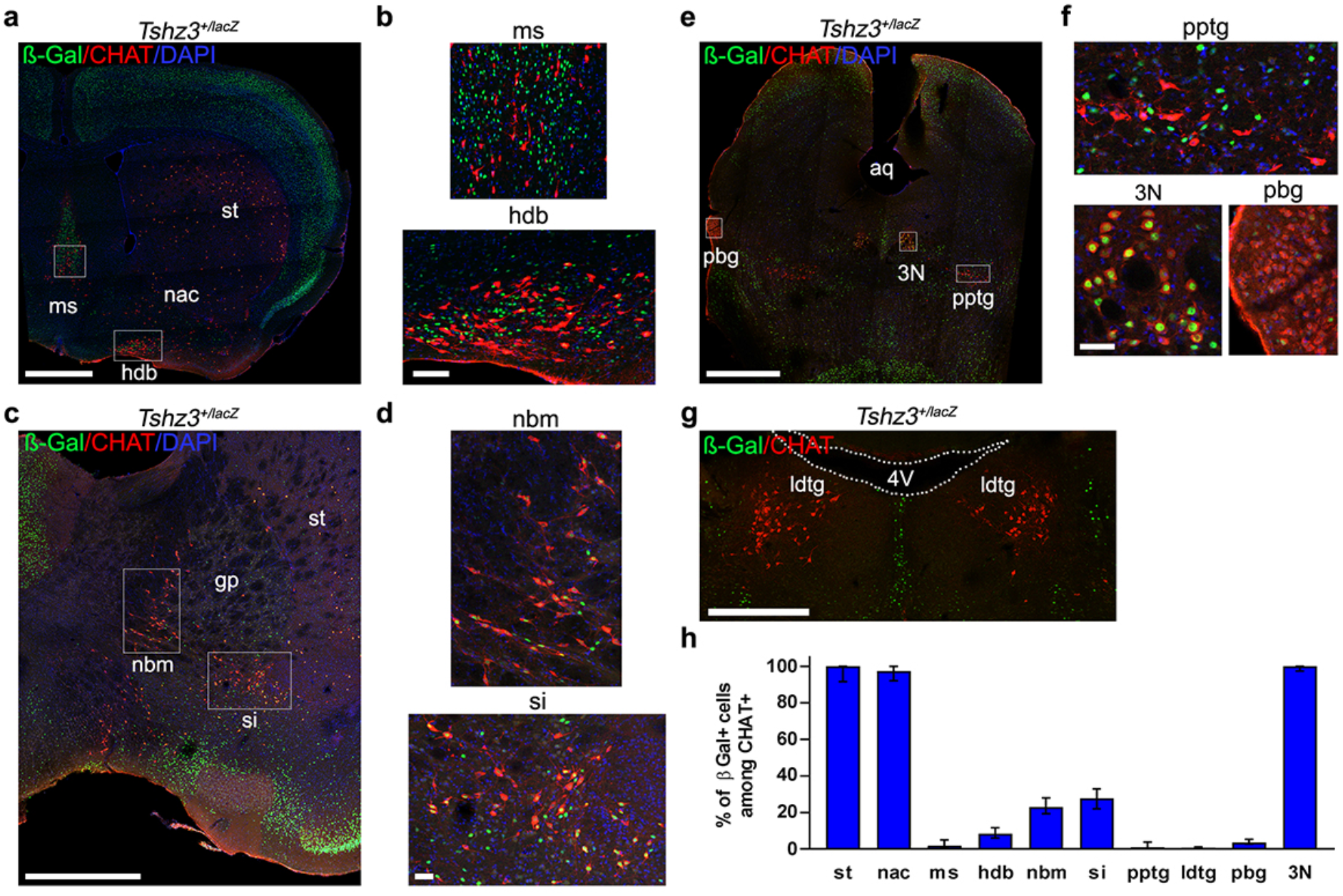
TSHZ3 expression in the main brain cholinergic systems. Forebrain (**a**-**d**) and brainstem (**e**-**g**) coronal sections stained for ß-Gal and CHAT. (**b, d, f**) Higher-power images of framed regions in **a, c** and **e**, respectively. **h** Quantification of ß-Gal-positive cells within the CHAT-positive population in brain structures containing cholinergic neurons. aq, aqueduct; hdb, nucleus of the horizontal limb of the diagonal band; gp, globus pallidus; ldtg, laterodorsal tegmental nucleus; ms, medial septal nucleus; nac, nucleus accumbens; nbm, nucleus basalis of Meynert; pbg, parabigeminal nucleus; pptg, pedunculopontine tegmental nucleus; si, substantia innominata; st, striatum; 3N, oculomotor nucleus; 4V, 4^th^ ventricle. Nuclei were counterstained with DAPI. Data are expressed as medians with interquartile range; they were obtained from 6 (3N), 7 (hdb), 9 (ms) 12 (pbg, si), 16 (ldtg), 17 (nac), 19 (st), 24 (pptg) and 40 (nbm) sections from 3 (hdb, ldtg, ms, pbg and pptg), 4 (si and 3N), 6 (nac), 7 (st) and 8 (nbm) mice, respectively.

**Figure S7.**
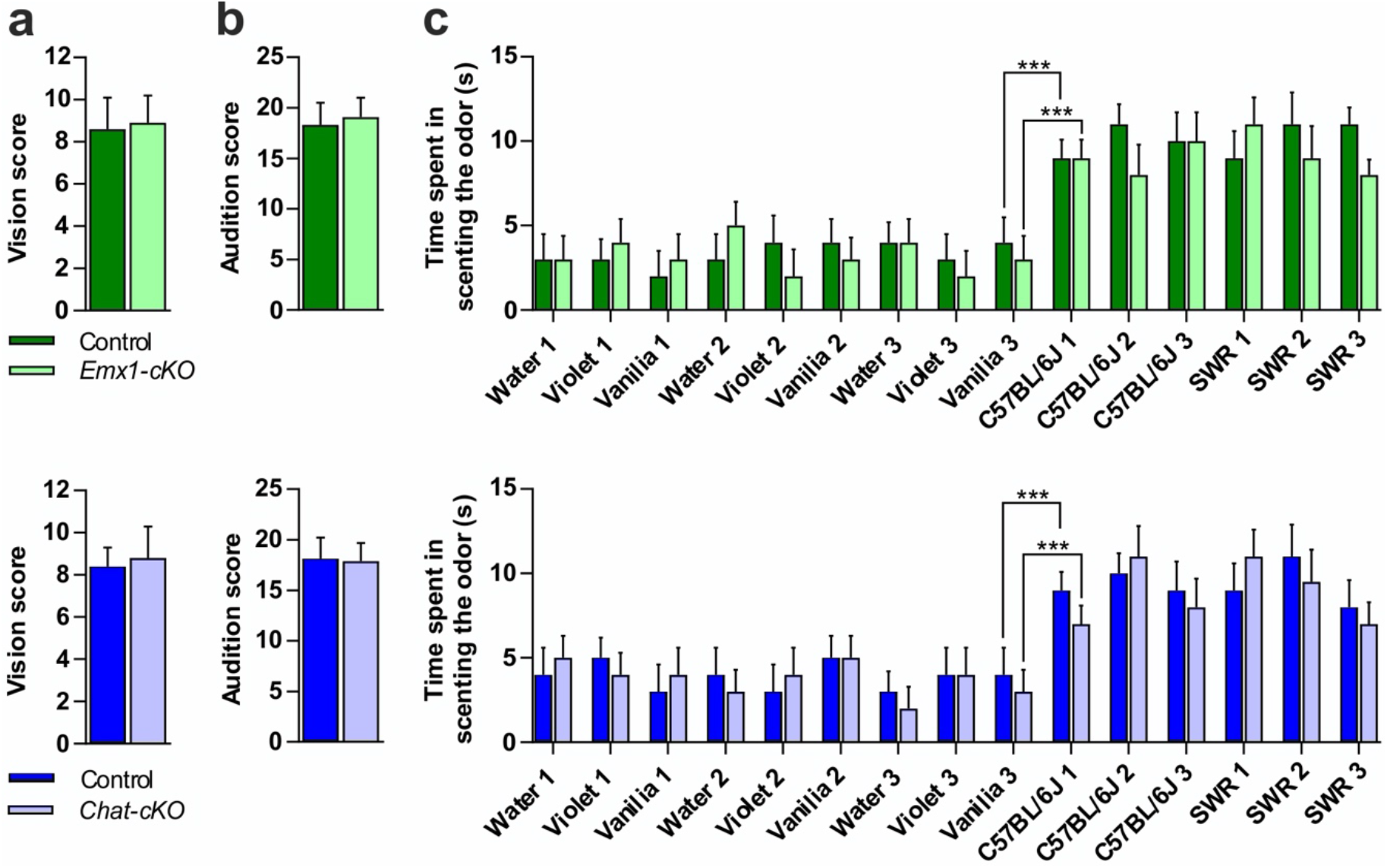
Visual, auditory and olfactory capacities in *Emx1-cKO* and *Chat-cKO* mice compared with their respective littermate controls. Ten mice per genotype were used in each screening. **a** Visual capacity differs neither in *Emx1-cKO* mice compared to their controls (Student’s *t* < 1, df = 18, non-significant (NS)), nor in *Chat-cKO* compared to their controls (Student’s *t* < 1, df = 18, NS). **b** Auditory capacities differ neither in *Emx1-cKO* mice compared to their controls (Student’s *t* = 1.2, df = 18, NS), nor in *Chat-cKO* mice compared to their controls (Student’s *t* < 1, df = 18, NS). **c** Time spent scenting non-social (water, violet, vanilla) and social (C57BL/6J, SWR) odors were analyzed with two mixed ANOVAs (*Emx1-cKO* and *Chat-cKO vs*. their respective control, and 15 odors as repeated measures). The genotype factor was not significant (*F* < 1, df = 1,18) in both cases. *Emx1-cKO, Chat-cKO* and their respective control spent more time sniffing social than non-social odors, as shown by comparing time sniffing vanilla 3 *vs*. C57BL/6J urine 1, the size of the differences being similar in each case for the KO and the control group (*Emx1-cKO* and control littermate: paired Student’s *t* = 4.5, df= 9, and *t* = 3.78, df = 9, respectively; *P* < 0.001; sizes of the differences : η ^2^ = 0.57 and 0.51, respectively; *Chat-cKO* and control littermate: paired Student’s *t* = 5.7, df = 9, and *t* = 4.9, df = 9, respectively; *P* < 0.001; sizes of the differences: η ^2^ = 0.49 and 0.40, respectively). Data are expressed as means + SEM. \*\*\**P* < 0.001.

**Figure S8.**
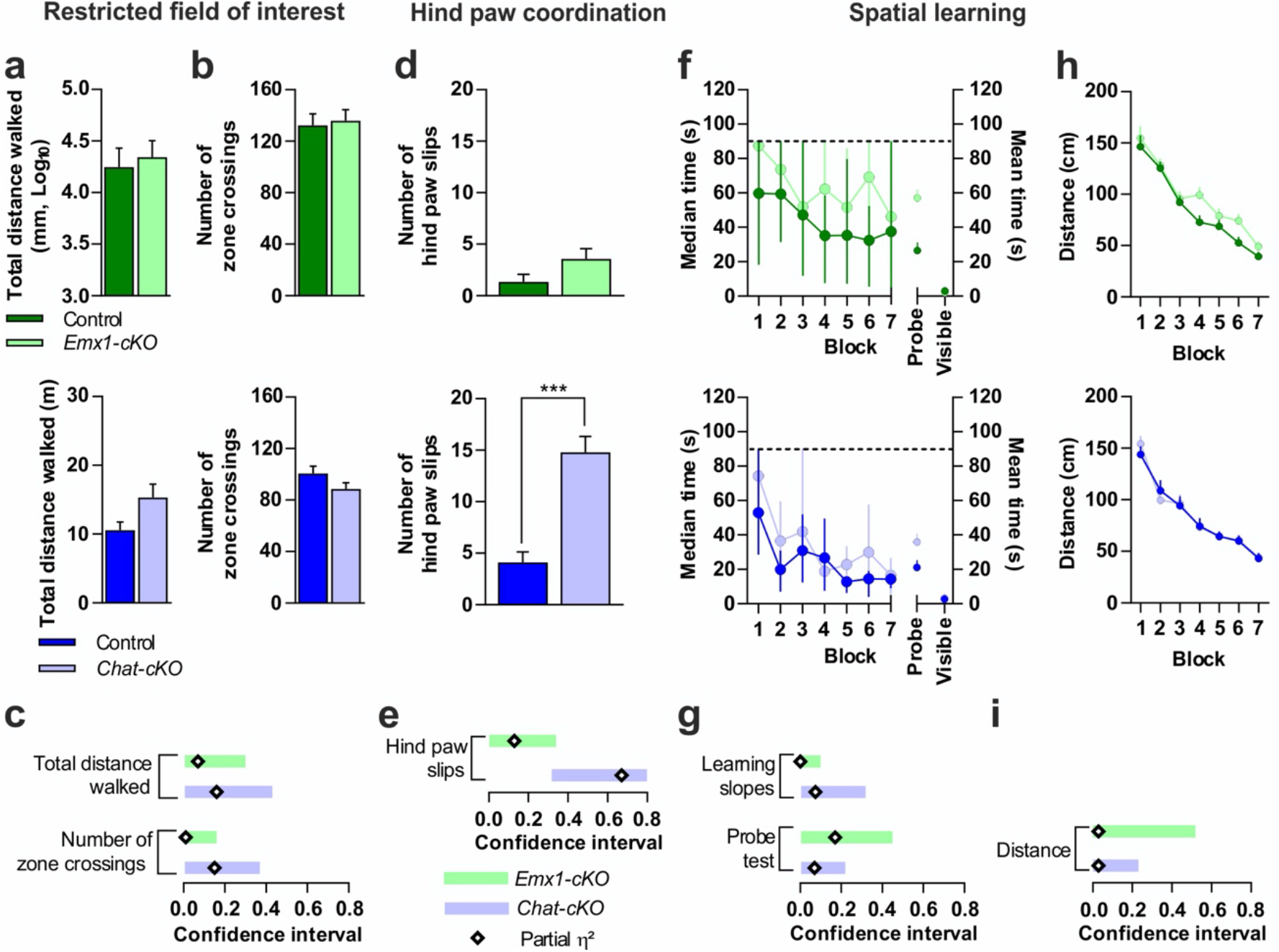
Restricted field of interest, hind paw coordination and spatial learning in *Emx1-cKO vs*. littermate control mice and *Chat-cKO vs*. littermate control mice. **a**-**c** The narrowness of the field of interest, expressed as the number of zone crossing in the open field **b** with the total distance walked **a** as covariate, is impacted neither in *Emx1-cKO* (n = 9) nor in *Chat-cKO* mice (n = 12) compared to their respective control (n = 8 and n = 8, respectively). **c** The partial η^2^ are very low and their confidence intervals includes zero. **d**-**e** Hind paw coordination. *Chat-cKO* mice (n = 9) exhibit a high deficit when compared to their control (n = 9) (Student’s *t* = 5.72, df = 16, P = 0.00003). On the opposite, *Emx1-cKO* mice (n = 10) do not differ from their control (n = 8) (Student’s *t* = 1.76, df = 16, *P* = 0.10). **e** The effect size of the difference in *Chat-cKO* (η^2^ = 0.67) exceeds the limit of impairment (0.30), whereas it is not considered in *Emx1-cKO* mice because its confidence interval encompassed zero. (**f**-**i**) Spatial learning in the Morris water maze. Time to reach the visible platform **f** is similar both in *Emx1-cKO* mice (n = 12) and their control (n = 11) and in *Chat-cKO* mice (n = 10) and their control (n = 13) (Student’s *t* = 0.90, df = 21, *P* = 0.38 and Student’s *t* = 1.28, df = 22, *P* = 0.21, respectively), showing that different learning performances cannot be attributed to motor or sensorial abilities. Non-parametric statistics were used in the hidden platform version when the assumption of normality of the distributions was rejected. We examined the learning slopes with the Friedman’s test for non-parametric ANOVA with repeated values. The four groups of mice learned across blocks 1 to 7. *Emx1-cKO* and their control learn equally (Friedman’s test for non-parametric ANOVA with repeated values: χ^2^ = 21.42, df = 6, *P* = 0.002 and χ^2^ = 19.22, df = 6, *P* = 0.004, respectively), with similar slopes (Student’s *t* = 0.01, df = 22, *P* = 0.99). *Chat-cKO* and their control also learned across blocks 1 to 7 with similar trends (χ^2^ = 24.41, df = 6, *P* = 0.0004 and χ^2^ = 30.67, df = 6, *P* = 0.00002, respectively) and similar slopes (Student’s *t* = 1.30, df = 21, *P* = 0.21). In the probe test version, the Student’s *t* in *Emx1-cKO vs*. control and *Chat-cKO vs*. controls are, respectively: Student’s *t* = 2.22, df = 22, *P* = 0.04 and Student’s *t* = 1.14, df = 21, *P* = 0.27. Dotted lines represent the 90 s cutoff. Dots indicating the visible platform values overlap. **g** The confidence intervals of the effect size for the learning slopes (η^2^ = 0.002 for *Emx1-cKO vs*. control and η^2^ = 0.07 for *Chat-cKO vs*. control) include zero, indicating that the difference of the learning slope can be disregarded. The confidence intervals of the effect size for the probe test (η^2^ = 0.17 for *Emx1-cKO vs*. control and η^2^ = 0.05 for *Chat-cKO v*s. controls) encompassed zero, indicating that the differences can be disregarded. **h** Cumulative distance from the hidden platform during the blocks. Learning was analyzed with parametric statistics (two-way mixed ANOVA with blocks as repeated-measures and cKO *vs*. control as between group variable). *Emx1-cKO* mice (n= 10) and their control (n= 12) learn equally (*F* = 63.18, df = 6,120, *P* = 7E-35, partial η^2^ = 0.76; interaction between blocks and groups (*F* < 1), with linear trend (*F* = 209.77, df = 1,20, *P* = 4E-12, partial η^2^ = 0.91)) and the slopes are identical (Student’s *t =* 0.76, df = 20, *P* = 0.46, η^2^ = 0.03). *Chat-cKO* mice (n = 10) and their control (n= 11) also learn equally (*F* = 71.44, df = 6,114, *P* = 2E-36, partial η^2^ = 0.79; interaction between blocks and groups (*F* < 1), with linear trend (*F* = 196.94, df = 1,19, *P* = 1E-11, partial η^2^ = 0.91)). The slopes are identical (Student’s *t* = 0.03, df = 19, P = 0.98, η^2^ = 0.00004). **i** The confidence intervals of the effect size for the learning slopes includes zero for both *Emx1-cKO* and *Chat-cKO vs*. their respective controls, indicating that the learning slopes do not differ in the two groups. Data are expressed as means + SEM (**a, b, d** and **h**), or as medians with interquartile range **f**. \*\*\**P* < 0.001.

